# Disturbance Sensitivity Shapes Patterns of Tree Species Distribution in Afrotropical Lowland Rainforests More Than Climate or Soil

**DOI:** 10.1101/823203

**Authors:** Chase L. Nuñez, James S. Clark, John R. Poulsen

## Abstract

Understanding how tropical forests respond to changes in the abiotic environment and human disturbance is critical for preserving biodiversity, mitigating climate change, and maintaining ecosystem services in the coming century. To evaluate the relative roles of the abiotic environment and disturbance on Afrotropical forest community composition we employ tree inventory data, remotely sensed historic climatic data, and soil nutrient data collected from 30 1-ha plots distributed across a large-scale observational experiment in previously logged, hunted, and pristine forests in northern Republic of Congo (Brazzaville). We show that Afrotropical plant communities are more sensitive to human disturbance than to climate, with particular sensitivities to hunting and distance to village (a proxy for other human activities, including tree-cutting, gathering, etc.). This study serves as an important counterpoint to work done in the Neotropics by providing contrasting predictions for Afrotropical forests with substantially different ecological, evolutionary, and anthropogenic histories.

## Introduction

Understanding how tropical forests respond to changes in human disturbance and changes in the abiotic environment is critical for preserving biodiversity, mitigating climate change, and maintaining ecosystem services in the coming century (Maslin *et al.* 2005; Malhi *et al.* 2013a). The lowland rainforests of Central Africa, in particular, are expected to lose 41% of their forest cover in by 2050 to forest clearing (Justice *et al.* 2001), largely due to expansion of subsistence agriculture and logging (Tyukavina *et al.* 2018). This loss in forest cover contributes to further tree mortality from drought and fire through indirect microclimatic shifts (Condit *et al.* 1996, 2017; Nepstad *et al.* 1999, 2004; Williamson *et al.* 2000; Martínez-Vilalta and Lloret 2016). Even modest changes in available water may push this area into a climatic space unviable for rainforests (Malhi and Wright 2004; Pan *et al.* 2011; Guan *et al.* 2015). Knowledge of how environmental disturbance has shaped current species and traits distributions could help more effectively allocate limited conservation resources to sensitive communities, yet our understanding is poor.

Current understanding of environmental effects on tropical forest communities comes largely from Neotropical studies that have focused primarily on the roles of drought and soil. In these drought-prone Neotropical environments, even short term water deficits induce important foliar adaptations in tropical forest trees that decrease leaf area and water usage (Nepstad 2002; Powell *et al.* 2017) despite having access to deep water reserves (Laskar and Robutel 2001; Broedel *et al.* 2017). Over time, low leaf area contributes to desiccation of leaf litter, and therefore can facilitate both fire (Nepstad *et al.* 2004) and tree mortality (Williamson *et al.* 2000; Martínez-Vilalta and Lloret 2016). This cascade of microclimatic change can then promote community turnover through the recruitment of shade intolerant species that are able to survive water-scarcity (Condit *et al.* 1995; Greenwood *et al.* 2017). Recruitment is also mediated by soil nutrient availability (Ceccon *et al.* 2003), contributing to a strong associations with tropical forest community composition (Plotkin *et al.* 2000; Harms *et al.* 2001) at local- (1 km^2^, John *et al.* 2007), meso- (1–100 km^2^, Clark *et al.* 1998, 1999) and landscape-scales (102 −104km^2^ ter Steege *et al.* 1993; Tuomisto *et al.* 1995; Phillips *et al.* 2003; Costigliola and Hogan 2016).

African tropical forests may respond differently to environmental changes than Neotropical forests due to community-level trait differences that are hypothesized to have arisen from Africa’s unique evolutionary past (Haffer 1969; Maley 1996; Maslin *et al.* 2005; Oslisly *et al.* 2013; Willis *et al.* 2013). Africa has fewer wet-affiliated species than would be expected from climate-biodiversity relationships (Leal 2009) and a high proportion of large trees that grow rapidly (Gond *et al.* 2013). Abnormally dry conditions during the last glacial maximum~ 26,500 years ago affected all tropical forests, yet only Afrotropical forests were reduced to small patches of remnant forest (Haffer 1969; Maley 1996; Maslin *et al.* 2005). These disturbances are hypothesized to have selected for species adapted to water scarcity and the capacity to disperse from refugia to recolonize landscapes cleared by receding glaciers (Leal 2009).

Paleoecological work shows evidence of Afrotropical forest communities’ resilience to past climatic and anthropogenic disturbances. Although forest composition was not completely unchanged (Brncic *et al.* 2007), moist semi-evergreen forest taxa have persisted throughout the last 3300 years with no sign of savannah expansion, despite evidence of anthropogenic activity and extensive periods of moisture limitation (Elenga *et al.* 2007). These studies based on pollen records cannot be extrapolated to current and future forest composition due to the uneven representation of species in sedimentary deposits (Mander and Punyasena 2018), so many researchers have turned to species distribution models (SDMs) that make use of recent presence-only, presence-absence, or abundance data to anticipate changes in forest diversity (Elith and Leathwick 2009). However, SDMs do not fully capture the degree to which species interactions mediate their response to the environment, contributing to a high level of uncertainty in predictions (Clark *et al.* 2017). This uncertainty is worsened by a reliance on parameters that have been fit at a simple level of aggregation (*e.g.* species-scale) to predict responses at more complex levels of aggregation (*e.g.* community-scale). The accumulation of error thorough aggregation is known as ‘Simpson’s Paradox’ or the ‘ecological fallacy’ (Bickel *et al.* 1975; Clark *et al.* 2011). Finally, SDMs have traditionally been taxonomically limited because they are not able to cohesively combine species data that are measured by different techniques and on different scales, or accommodate data with most species are absent from most sites (Clark 2016; Clark *et al.* 2017; Taylor-Rodríguez *et al.* 2017). Improved predictions can be obtained by using a model, like the Generalized Joint Attribute Model (GJAM), that incorporates information about the associations between species to determine the joint community response (Clark *et al.* 2017).

To evaluate the relative roles of the abiotic environment and disturbance on Afrotropical forest community composition we employ tree inventory data, remotely sensed historic climatic data, and soil nutrient data collected from 30 1-ha plots distributed across a large-scale observational experiment in previously logged, hunted, and pristine forests in northern Republic of Congo (Brazzaville). We then use plant trait data derived from the literature to explore the potential role of plant functional traits on community composition. This is the first experiment to our knowledge that directly tests the relative effects of competing drivers of community structure and composition using an integrative data-fusion approach. We hypothesize that species will be more sensitive to human disturbance relative to climate and soil (Haffer 1969; Maley 1996; Maslin et al. 2005; Oslisly et al. 2013; Willis et al. 2013). Our null assumption is that these communities will mirror findings in Neotropical forests and show the greatest relative sensitivity to dry season precipitation and temperatures (Engelbrecht et al. 2007), followed by soil nutrients (Cook *et al.* 1992; Tuomisto *et al.* 1995; Clark *et al.* 1998; Plotkin *et al.* 2000; Harms *et al.* 2001; Phillips *et al.* 2003; John *et al.* 2007; Costigliola and Hogan 2016) and human disturbance (Peres et al. 2010; Solar et al. 2016).

## 1.2 Methods

### Study Area

We conducted the study in the Nouabale Ndoki National Park (NNNP; 400,000 ha) and the Kabo logging concession (267,000 ha) in northern Republic of Congo (Figure 1). The forests in this area are classified as lowland tropical forest where *Meliaceae, Euphorbiaceae*, and *Annonaceae* are the dominant tree species (CIB 2006). This area receives an average rainfall of about 1700 mm annually with seasonal peaks in May and October. The Kabo concession borders the NNNP to the south, and together they form a mosaic of logged and unlogged forest. Twenty years before the study began in 2005, the Kabo logging concession was selectively logged at low intensity (<2.5 stem ha^−1^) with four species making up 90% of the harvest volume: *Entandophragma cylindricum*, *E. utile*, *Triplochiton scleroxylon*, and *Milicia excelsa* (CIB 2006). Approximately 3000 people inhabited the study site at the time of the study, most residing in the logging town of Kabo. Residents mainly hunt with shotguns, and occasionally with wire snares, for consumption and for local trade (Poulsen *et al.* 2009). A gradient of hunting intensity decreases with distance from Kabo (Poulsen *et al.* 2011), with some forest types being used more than others (Mockrin 2008). This gradient is captured best in the Euclidian distance predictor variable ‘distance to nearest village’ because it shows the manifold types of human uses of forests that dissipate as human activities become less intense.

**Figure 1:**
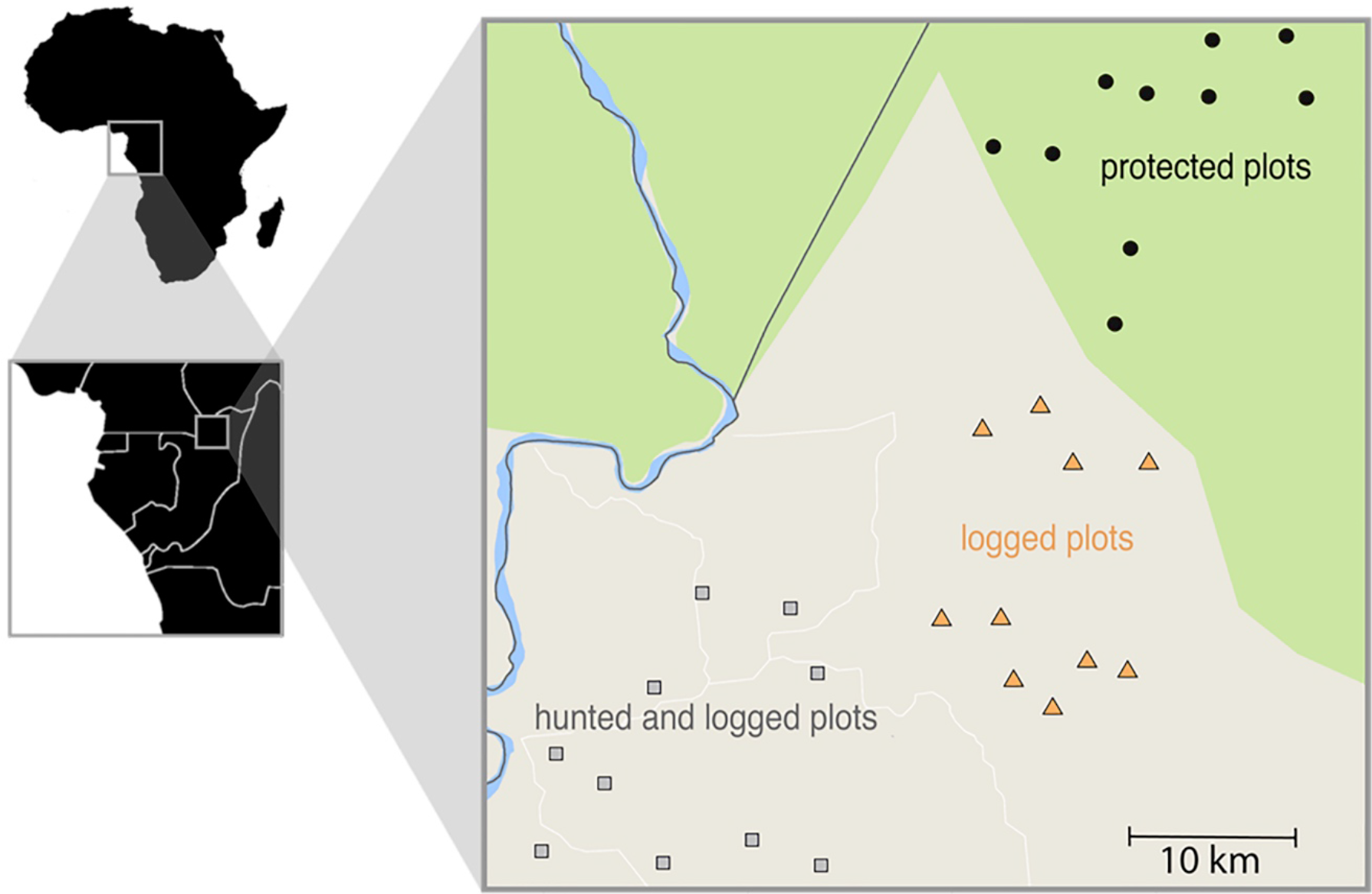
Location of 30 1-hectare study plots in Central African Country of Republic of Congo (Brazzaville). Plots were situated in forest that were unlogged and unhunted (10 plots, circles), sites that were logged and unhunted (10 plots, triangles), and sites that were both logged and hunted (10 plots, squares). Protected plots fall within the border of Nouabalé -Ndoki National Park whereas plots exposed to hunting and/or logging were located in the Kabo logging concession.

### Tree Census Data

We established 30 1-ha tree plots; 10 sites were situated in forest that were unlogged and unhunted, 10 sites that were logged and unhunted, and 10 sites that were both logged and hunted. Using ArcView 3.2 and a 14 class habitat map (Laporte et al. 2007), we randomly located plots within each disturbance regime in mixed lowland forest, with a buffer of at least 500 m to the nearest primary road and 100 m to the nearest water source. Within each plot, all trees greater than 10 cm diameter at breast height (DBH) were tagged, measured, mapped, and identified to species (Wortley and Harris 2014). We additionally recorded canopy status (understory, mid-story, canopy, and emergent), canopy openness, and light availability by averaging values from four hemispherical pictures taken at each quarter of a plot.

### Plant Species Trait Data

Life history traits for the tree species found in the 30 1-ha plots were derived from the TRY global database of curated plant traits (Kattge *et al.* 2011). This included seed mass (Kühn *et al.* 2004), wood density (Zanne *et al.* 2009) and foliar traits (Kirkup *et al.* 2005; Kerkhoff *et al.* 2006; Kraft *et al.* 2008; Powers and Tiffin 2010; Vergutz *et al.* 2012; Meng *et al.* 2015). Trait values were taken for every species available (< 1% of species). If species level data were not found (~ 21%), we used the genus average. If neither genus nor species level were available (~ 78%), we used the family average. It would be ideal to have trait mean and variance information for all species, but this is not possible in understudied areas with high biodiversity. This method of phylogenetic trait interpolation, although rudimentary, allows us to use the best available information while not biasing the analysis. That is, species with less phylogenetically specific information will be supplied with values approaching the mean, and therefore have no predictive power on the response.

### Soil Data

We collected soil samples from the corners and center of each plot using a soil probe (2.85 3 83 cm) to 15 cm depth. Samples were weighed (wet mass), and then air-dried and weighed again (dry mass). For analysis, the five samples from each plot were combined and homogenized before being sent to the IFAS Extension Soil Testing Laboratory at the University of Florida. Homogenized soil samples were analyzed for percentage sand, clay, and silt, as well as pH and nutrient availability (N, P, K, Al, Ca, Mg, Mn). Available cations and P were extracted using the Mehlich III solution (Tran and Simard 1993). Elemental analysis for the cations and P was done on the extracts by using inductively coupled plasma (ICP) spectroscopy. We extracted N as NH4 and NO3. Nitrogen was estimated calorimetrically using a Technicon II auto-analyzer (SEAL Analytical, Mequon, Wisconsin, USA). The Kjeldahl method was used for the determination of total N (Hesse 1971). Soil pH was measured in an Adams-Evans buffer solution made up of one volume of soil diluted in two volumes of water. Subsoil samples were analyzed for soil texture, using the hydrometer method (Sheldrick and Wang 1993).

### Climate Data

Average historical precipitation, potential evapotranspiration (PET), maximum temperature, and minimum temperature for each plot were derived from the NASA TerraClimate product (Abatzoglou et al. 2018) for 1985 – 2017 accessed using Google Earth Engine (Gorelick et al. 2017). To capture the contribution of seasonality, we amalgamated historical climate data before model selection. Climate values were averaged within the three recognized seasons in this region: long rains: (May - July), short rains (September - October), and dry season (November - April).

### Generalized Joint Attribute Model for Species Distribution

We employ a generative Generalized Joint Attribute Model (GJAM) that predicts species abundance at the scale and context used to fit the model: jointly, on the community scale (Clark *et al.* 2017). GJAM estimates can therefore be interpreted on the scale of the observations, accounting for sample effort. The model is based on a joint distribution [θ,X,Y] of parameters θ, predictors X, and species responses Y. Parameters (θ) in the model include matrices of coefficients relating X to Y (B) and the residual covariance matrix for all species pairs in Y (Σ). In effect, Σ represents the covariance between species beyond what has already been explained by the environmental covariates. It can include interactions between species, unaccounted for environmental gradients, and other unexplained sources of error. In the generative model, all elements, including estimates and prediction, are part of one analysis. The likelihood is: [Y_1_,…,Y_S_ |θ,X], where subscripts refer to species 1 through S. Model fitting is done on the shared prediction/observation scale, based on the posterior distribution, [θ |X,Y]∞[Y1,…,YS |θ,X][θ]. On the right-hand side is the likelihood and the prior distribution, [θ], which is non-informative. Sensitivity of the entire response matrix to environmental predictors can be obtained from the diagonal vector of the covariance matrix between predictors in X in terms of the responses they elicit from Y.

### Generalized Joint Attribute Model for Traits

The incorporation of plant traits in GJAM is consistent with the structure of the model listed above, with community weighted mean (CWM) trait composition being predicted instead of species counts – *i.e.* the response is a trait by species matrix Y, which could be any trait or traits of interest. Use of CWM traits, or trait values weighted by the number of species in a plot, is a useful method of broadly characterizing communities, illuminating relationships between environmental gradients and patterns of life history adaptations and community assembly through environmental filtering (Wright *et al.* 2004; Cornwell and Ackerly 2009; Sonnier *et al.* 2010). However, because trait data come in many forms (*e.g.* categorical classes, continuous measurements, discrete counts, etc.), GJAM leverages censoring with the effort for an observation to combine continuous and discrete variables with appropriate weight. In count data, effort is determined by the size of the sample plot, search time, or both. It is comparable to the offset in GLMs. Full model specifications can be found in (Clark *et al.* 2017). The suite of covariates (Appendix, A1) included in the final model were selected by iteratively combining each covariate and comparing the Deviance Information Criterion (DIC) and Variance Inflation Factor (VIF) values for each model structure (including interactions) and selecting the optimal model that produced the most inference on the relationship between predictors and responses, while returning low DIC and VIF less than 3. Model estimates were taken from 100,000 iterations, discarding the first 1000 iteration as pre-convergence. We visually inspected trace plots to confirm convergence and adequate mixing (Appendix A6).

## 1.3 Results

Responses to the environment, *i.e.* the effect of the environment on species counts, varied among species (Appendix A2), but covaried within two groups with opposing responses to predictor variables (Figure 2B). Group 1 (disturbance tolerant species, Figure 2B1) was more likely to occur in plots disturbed by hunting, logging, or both, while group 2 (disturbance intolerant species, Figure 2B2) was most likely to occur in pristine plots undisturbed by hunting or logging. The varied responses of all species can be summarized by a comparison of community sensitivity to predictors across all species (Figure 3, S5), with greatest sensitivity (*i.e.* the relative increase in predictor to produce a one unit increase in species counts) to human disturbance (distance to nearest village 3.29 [2.76, 3.85], hunting 2.22 [1.85, 2.61], and logging 1.99 [1.65, 2.35]), followed by moderate sensitivity to dry season temperature (dry season minimum temperature 1.48 [1.21, 1.78], dry season maximum temperature 1.28 [1.06, 1.51]), and least sensitivity to dry season precipitation (0.92 [0.76,1.09]) and soil composition TKN (1.15 [0.94, 1.38]); P (1.11 [0.92, 1.31]); K (1.08 [0.91, 1.27]); pH (1.02 [0.85, 1.20]).

**Figure 2:**
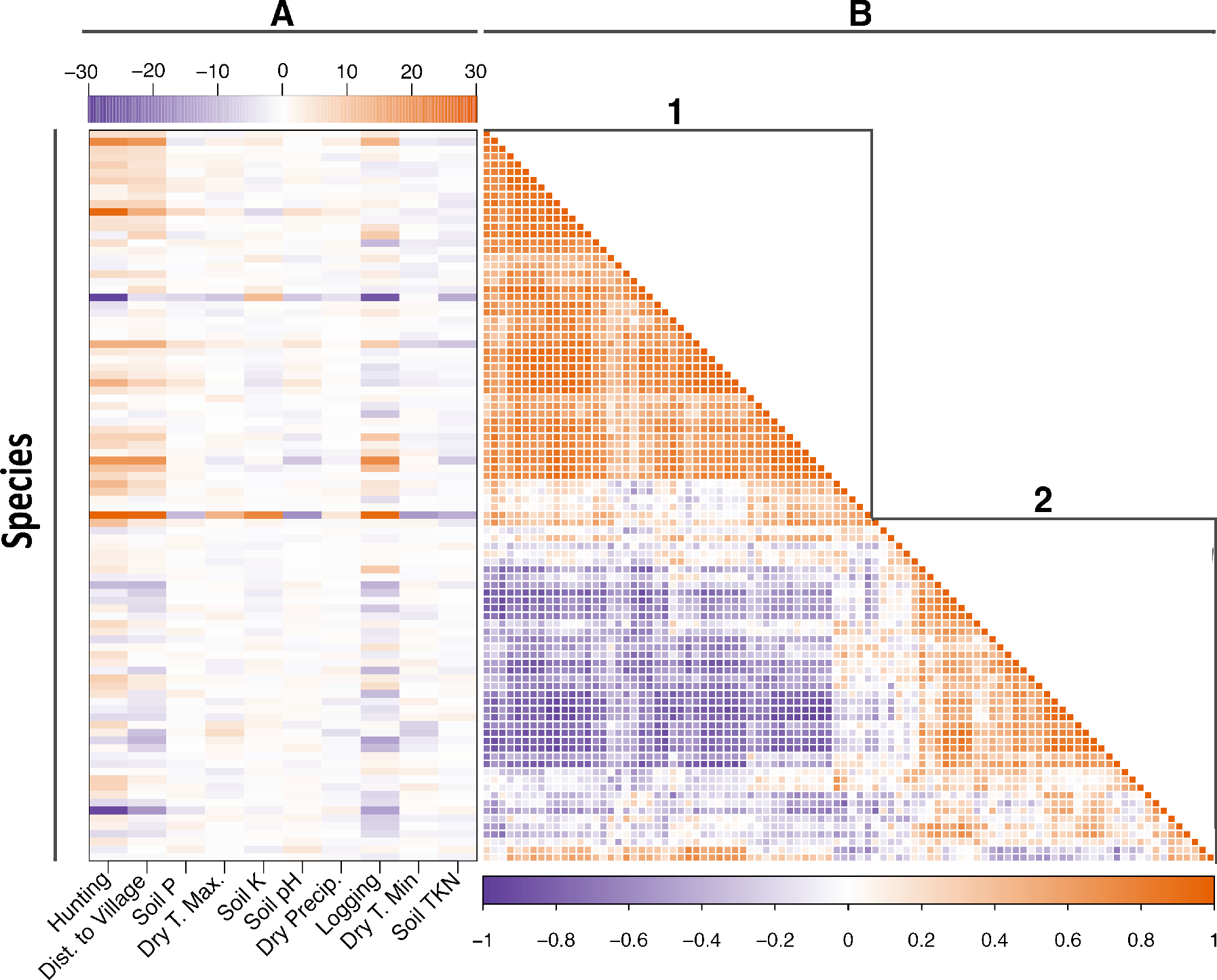
Posterior parameter estimates for the effect of environmental covariates on species counts (A) and covariance between species (B) show clustering into two general groups that represent a disturbance tolerant pioneer community (B1) and disturbance intolerant species (B2). Species names are listed in Appendix A9.

**Figure 3:**
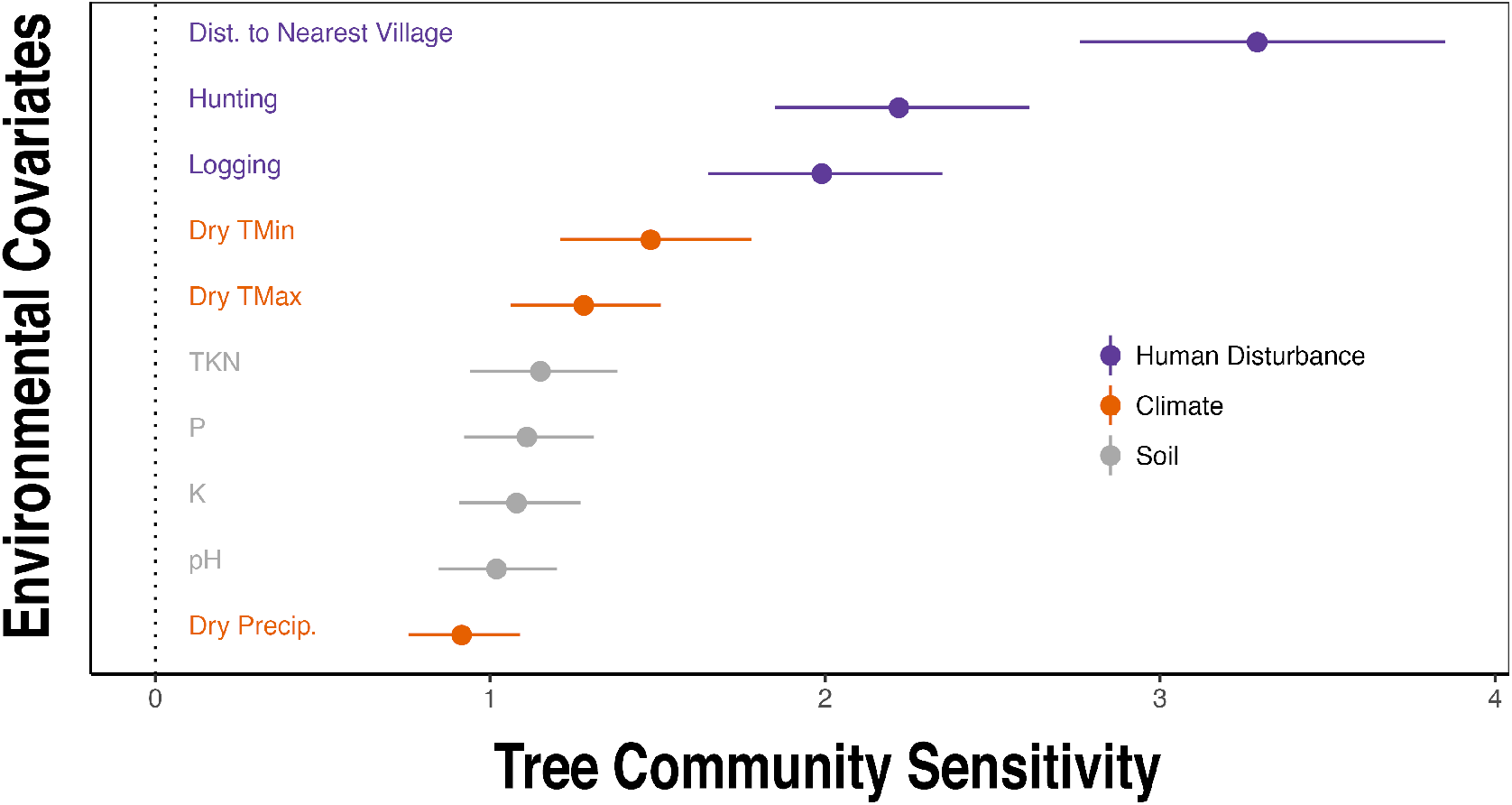
Sensitivity of community species composition to environmental predictors and 95% credible interval (CI). Sensitivity is dimensionless and in the unit of the environmental covariate.

Our trait analysis indicates that soil pH was strongly related to all traits except sapwood density (S6). Soil nutrients were weakly informative of community weighted trait values: total nitrogen (TKN) was weakly related to leaf N (1.35 × 10^−06^ [2.70 × 10^−07^, 2.41 × 10^−06^]), and N15 (6.1 × 10^−04^ [9.62 × 10^−05^, 1.12 × 10^−03^]), as well as C13: −1.27 × 10^−03^ [−1.74 × 10^−03^, −8.04 × 10^−04^]). Soil P was moderately informative of the distribution of species with high leaf C:N ratios (0.20 [0.02,0.38]), C13 (−0.18 [−0.24,−0.12]), and leaf surface area (0.04 [0.00,0.07]). Soil K was weakly informative of species with greater leaf C13 (0.02 [0.01,0.03]).

There was a strong relationship between dry season climate and community weighted leaf traits: maximum temperature on chlorophyll concentration (2.00 [1.37, 2.65]), carbon to nitrogen ratio (0.52 [0.25, 0.78]), surface area (0.11 [0.06, 0.16]), toughness (0.04 [0.00, 0.07]), sapwood density (0.02 [0.01, 0.03]), C13 (−0.84 [−0.93, −0.75]), and SLA (0.48 [0.28, 0.68]) (Figure 4A, S6).

**Figure 4:**
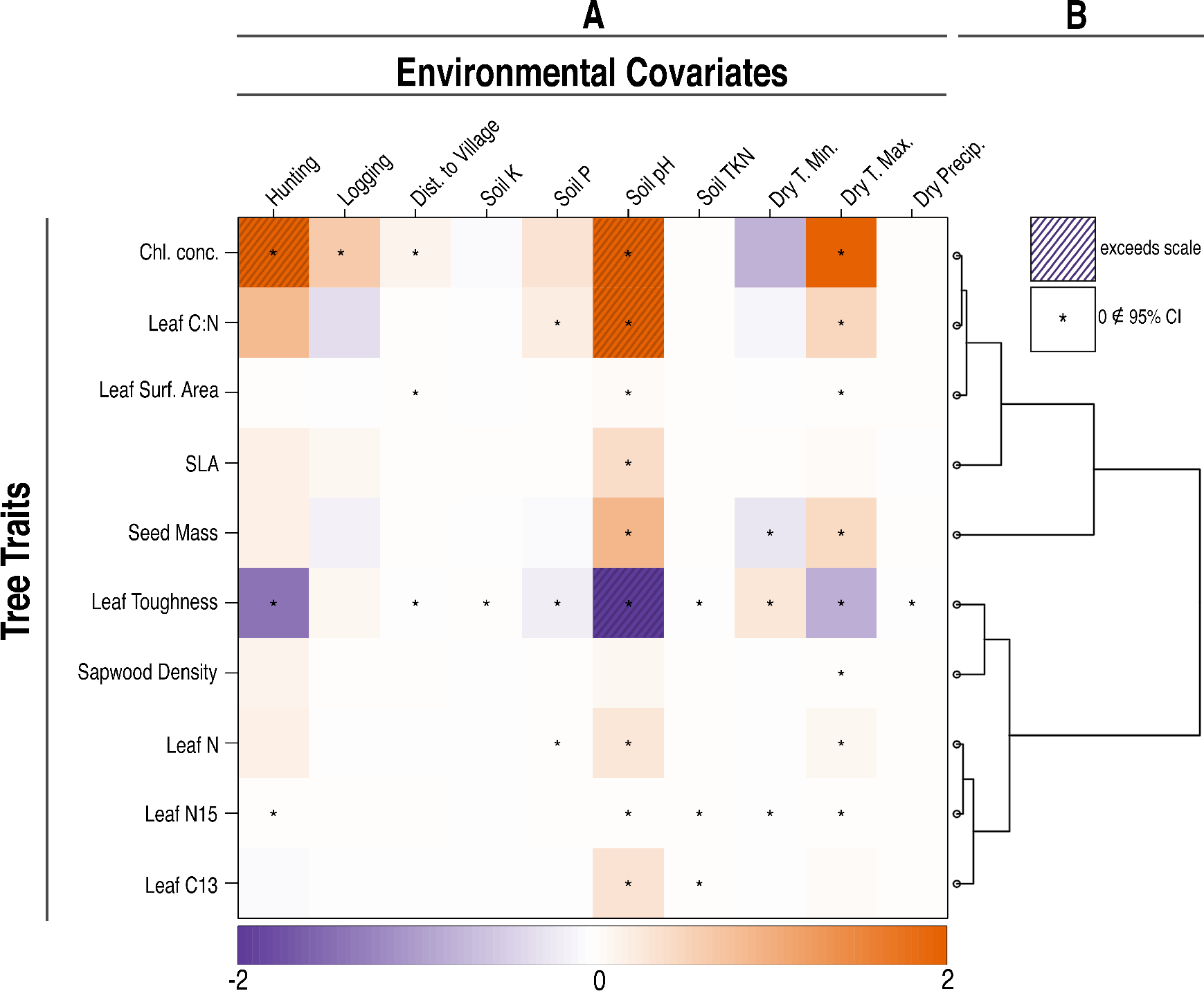
Standardized posterior parameter estimates for effect of climatic and human disturbance predictors on community weighted traits (A) and a dendrogram of correlation in trait response to predictors (B).

Disturbance was strongly related to just two specific traits: chlorophyll concentrations (hunting (3.75 [0.85, 6.65]), distance to nearest village (0.16 [0.05, 0.28])) and leaf C13 (hunting (−1.42 [−1.82, −1.02]), and distance to nearest village (−0.03 [−0.05, −0.02])).

Community weighted traits covaried in two groups: one encompassing the similar responses of chlorophyll concentration, leaf C:N, leaf surface area, SLA, and seed mass to environmental predictors, and the other encompassing the similar responses of leaf toughness, sapwood density, and leaf N, N15, and C13 (Figure 4B). Community weighted traits were most sensitive to human disturbance (distance to nearest village 17900 [793, 48100], hunting 10300 [437, 30200], and logging 10500 [447, 30200]); followed by moderate sensitivity to soil composition TKN (6650 [289, 18300]), P (9920 [499, 25700]), K (6430 [268, 18400]), pH (8700 [419, 22100]), and dry season climate (dry season minimum temperature 8370 [360, 24300], dry season maximum temperature 6450 [274, 18900], and dry season precipitation 4720 [198, 13800]) (Figure 5, S7).

**Figure 5:**
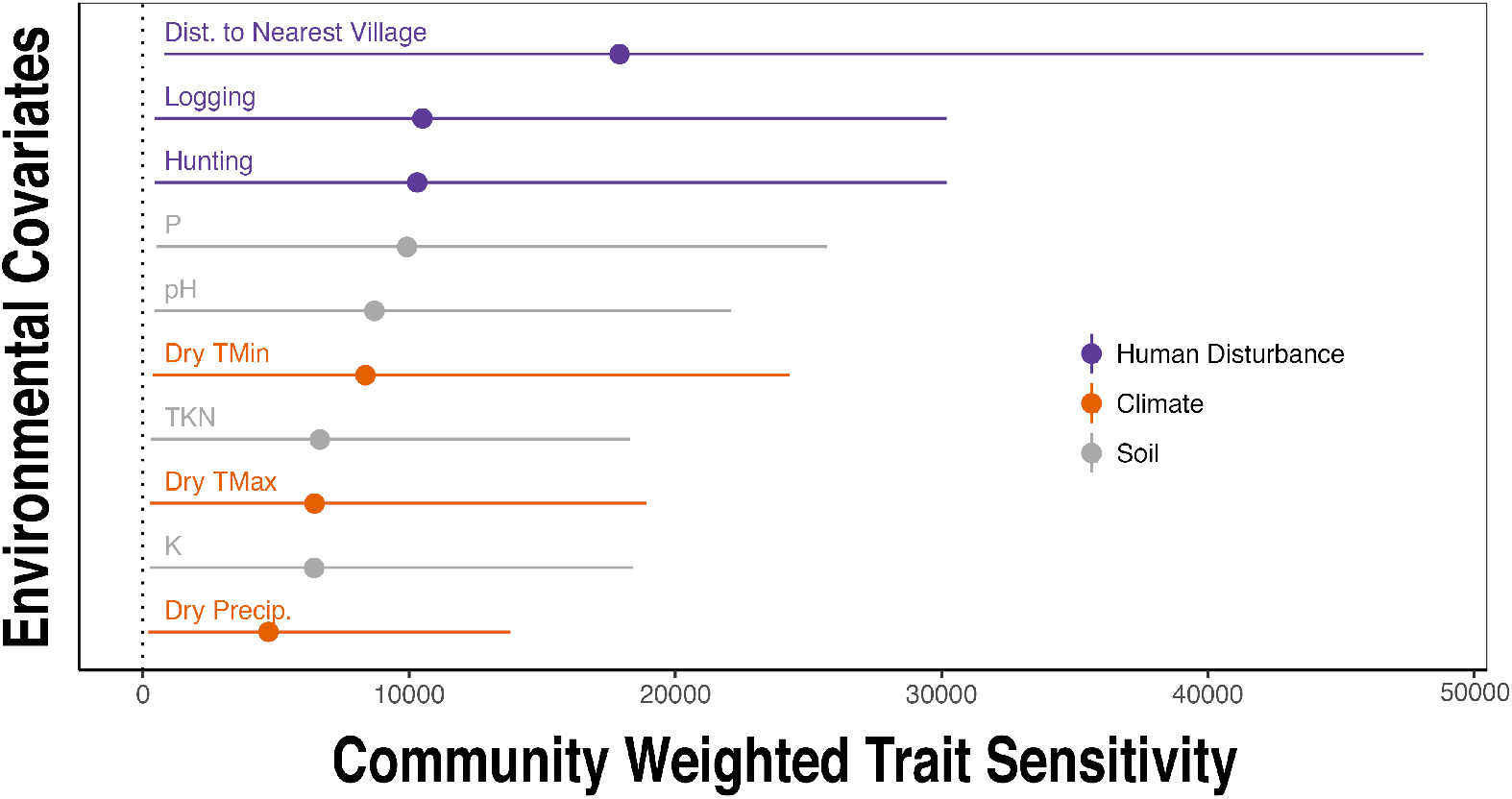
Sensitivity of community weighted traits to environmental covariates and 95% credible interval (CI). Sensitivity is dimensionless and in the unit of the environmental covariate.

## 1.4 Discussion

We find Afrotropical plant communities are more sensitive to human disturbance than to climate or soil. This result contrasts with Neotropical studies that show strong sensitivity to dry season precipitation and temperatures (Engelbrecht et al. 2007), and soil nutrients (Cook *et al.* 1992; Tuomisto *et al.* 1995; Clark *et al.* 1998; Plotkin *et al.* 2000; Harms *et al.* 2001; Phillips *et al.* 2003; John *et al.* 2007; Costigliola and Hogan 2016). We show that responses to the abiotic environment varied among species (Figure 2A, S4), but covaried within two groups with opposing responses to predictor variables (Figure 2B), particularly hunting and distance to village (an inclusive proxy for other human activity) that disproportionately affect species composition in these plots (Figure 3, S5). Human activity has well known negative effects on Central African animal communities (Poulsen *et al.* 2011, 2017; Koerner *et al.* 2017), and our results suggest that these defaunation gradients also have long-lasting effects on whole ecosystems through indirect stress on plants and alterations in plant-animal interactions that can modify plant community composition (Poulsen *et al.* 2013; Nuñez *et al.* 2019).

Our trait (Figure 4A, S6) and sensitivity analyses (Figure 5, S7) demonstrate that this grouped response of species to human disturbance is likely being driven by traits associated with two distinct successional communities. Human disturbance predictors were strongly related to a subset of foliar traits common to disturbance related pioneer species – higher chlorophyll concentrations and leaf surface area, with lower leaf toughness – as well as moderate sensitivity to soil composition and dry season climate. The considerable uncertainty surrounding estimates of predictors on community weighted traits is likely due in part to our coarse trait data (see below), but could also indicate that these communities are not being assembled through systematic trait-filtering, but rather a species-specific response to both measured and unmeasured variables in ways that are not readily decomposed into traits (Clark 2016).

Contrary to past studies finding a strong effect of soil composition on species composition (Cook *et al.* 1992; Tuomisto *et al.* 1995; Clark *et al.* 1998; Plotkin *et al.* 2000; Harms *et al.* 2001; Phillips *et al.* 2003; John *et al.* 2007; Costigliola and Hogan 2016), our results indicate that soil heterogeneity had weaker effects on tree communities than other variables (Figure 3). Soil characteristics were more strongly related to community weighted leaf traits in accordance with ecophysiological theory (Vitousek 1984; Ordoñez *et al.* 2009; Mayor *et al.* 2014), including a particularly strong relationship between soil pH and all plant traits except sapwood density (Figure 4A, S6). This is likely because soil pH indirectly affects the uptake and availability of several plant nutrients. Ca, Mg, K, and P are less available in low pH soils, whereas Al, Cu, Mn, and Zn cations become more available in low pH soils (John *et al.* 2007; Daniels 2016). Nevertheless, it is important to consider tropical forest soil dynamics when interpreting these results. Tropical soils are notoriously nutrient poor as a result of most nutrient matter being held in the aboveground biomass; however, it is likely that the observed soil nutrient composition at the time of study does not accurately represent the nutrient availability when the current adult trees were dispersed as seeds and recruited into the canopy.

Climate had moderate effects on community composition as species abundances were most sensitive to dry season temperature and to some degree sensitive to dry season precipitation (Figure 3, S5). There was also a strong relationship between dry season climate and community weighted leaf traits (Figure 4A, S6), likely due to the role those traits have in moderating evapotranspiration and maximizing photosynthesis. These effects, although slight in comparison to human disturbance, could have large effects on forest composition in light of the rapid increases in temperature predicted for this area (Leal 2009; Malhi *et al.* 2013b)—up to a 6°C increase by end of century in the high emission scenario (CSC 2011).

Care needs to be taken when interpreting results from tree plots. Although individual species locations and identities in our dataset are quite refined, environmental predictor data (minimum/maximum temperature, precipitation) are spatially coarse. The spatial extent of our study design covers ~4000 km^2^ and three disturbance regimes, but there is still relatively little variation in climate at this scale, potentially reducing their predictive power. Additionally, strong effects of human disturbance could be partially due to our study design, which was necessarily pseudoreplicated, *i.e.* study plots affected by the same disturbance type were geographically grouped together because human disturbance was also geographically concentrated. This was a direct result of the spatial pattern of hunting and logging around the village of Kabo (Poulsen *et al.* 2011), and means that other, unmeasured environmental gradients could have influenced our results. Future work should be conducted at a larger spatial extent (*e.g.* nationally with independent replicates for each disturbance regime) to see if the effects of disturbance hold up at larger spatial scale. Additionally, interpolated trait data require careful *post hoc* interpretation. Trait data do not exist for the majority of tropical species, and therefore values used are community averages for each plot calculated from the low proportion of species for which data were available. To model future responses of tree species and communities to changing climate, we need higher resolution species trait and local weather data, both lacking in developing tropical areas. Effort and investment should be committed to collecting trait data for tropical tree species for upload onto open databases like TRY (Kattge *et al.* 2011), as well as for increasing the number of local weather stations across Central Africa.

Our findings represent the most thorough study so far linking the joint responses of Afrotropical tree species distribution patterns with species’ environmental responses. The response of tropical plant species to environmental gradients like soil nutrients, light, and water have been experimentally measured in the past (Hill and Hamer 2004; Palmiotto *et al.* 2004; Ewel and Mazzarino 2008; Clark *et al.* 2016; Martínez-Vilalta and Lloret 2016), but how these responses control whole communities has been less well understood. This is the first study to our knowledge to jointly model the response of Afrotropical forest communities to soil, climate, and disturbance. Our results emphasize the sensitivity of tropical forests to both human activity and climate, with human disturbance having a direct role in determining species distributions in concert with local and regional water availability. Thus, the future expansion of humans into Central African forests coupled with dramatic increases in temperature expected by the end of the century will have direct consequences for species ranges, tropical forest community composition and ecosystem function. Conservation of tree communities will necessary involve mitigating disturbances of hunting and logging in the short-term and climate change in the long-term. Current vegetation–climate models, particularly for tropical regions, suffer from a lack of ecological data and mechanistic understanding of the factors shaping current species distributions. The knowledge that dry season temperature, together with the sensitivity of species to human disturbance, is influencing species distribution patterns in tropical forests will help to improve the accuracy and specificity of predictions of vegetation shifts under global change scenarios

