## Appendix A for "Disturbance Sensitivity Shapes Patterns of Tree Species Distribution in Afrotropical Lowland Rainforests More Than Climate or Soil"

**A1: Comparison of scaled covariates across 30 study plots shows greatest climatic variation across plots in dry season and heterogeneous plot-level soil characteristics.**

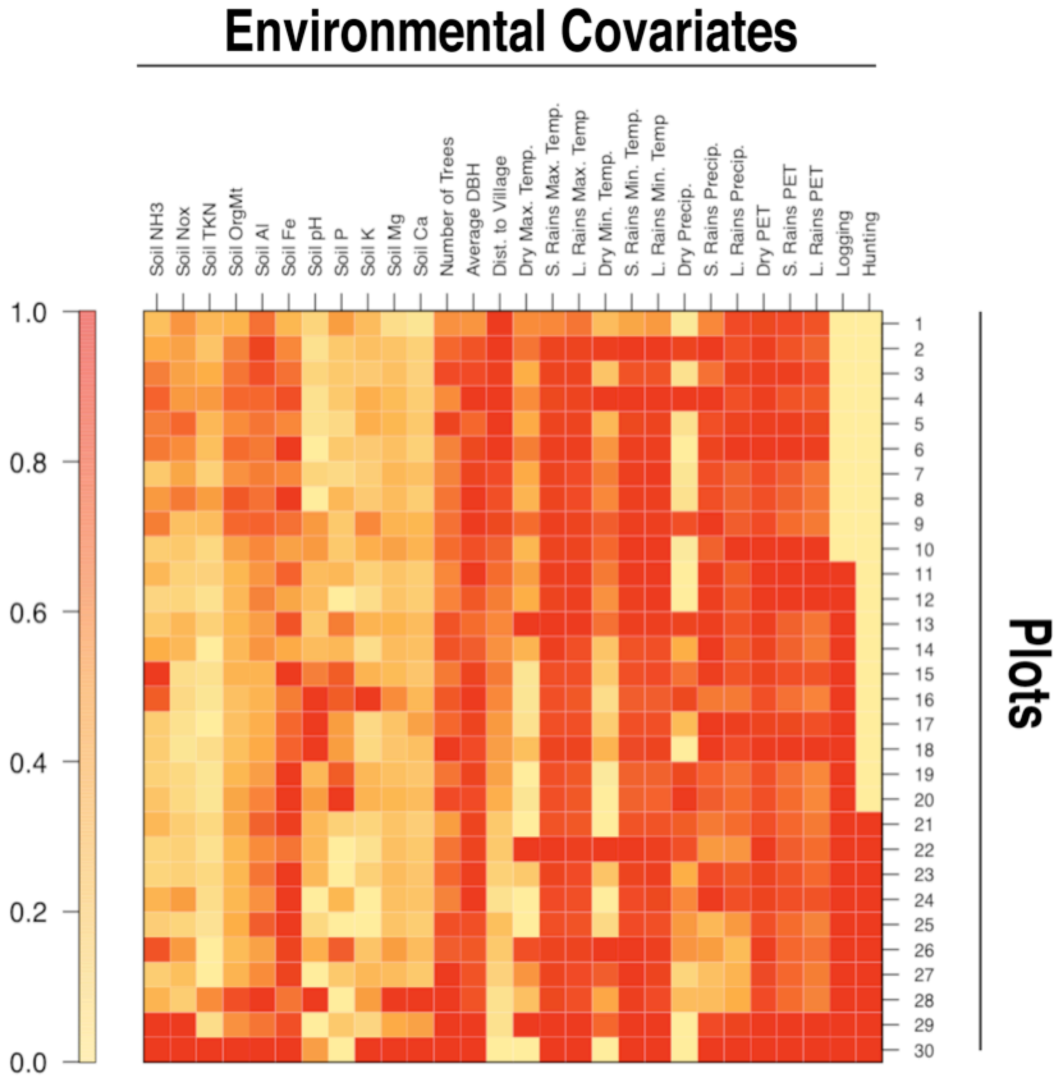

**A2: Species-specific posterior parameter estimates for the effect of each environmental covariate on species counts.**

Posterior Parameter Estimates for Species

| | Estimate | 2.50% | 97.50% | 0 ∉ 95% CI |
|---|---|---|---|---|
| Afrostyrax lepidophyllus | | | | |
| intercept | 0.77 | -0.04 | 1.62 | |
| hunting1 | 6.63 | -3.85 | 17.20 | |
| logging1 | -3.41 | -13.00 | 6.06 | |
| Dist | 0.24 | -0.16 | 0.64 | |
| K | -0.18 | -0.35 | -0.02 | * |
| P | 1.27 | -0.35 | 2.93 | |
| Ph | 5.17 | -2.42 | 13.10 | |
| TKN | 0.00 | -0.02 | 0.01 | |
| tmin.dry.mean | -1.75 | -4.79 | 1.15 | |
| pr.dry.mean | 0.02 | -0.04 | 0.08 | |
| tmax.dry.mean | 0.68 | -1.52 | 2.93 | |
| Afzelia bipindensis | | | | |
| intercept | 0.44 | -0.34 | 1.22 | |
| hunting1 | 3.35 | -6.08 | 12.80 | |
| logging1 | -0.92 | -9.81 | 7.88 | |
| Dist | 0.04 | -0.33 | 0.41 | |
| K | 0.01 | -0.14 | 0.16 | |
| P | 0.32 | -1.11 | 1.73 | |
| Ph | 4.09 | -2.68 | 10.90 | |
| TKN | 0.00 | -0.02 | 0.01 | |
| tmin.dry.mean | 0.96 | -1.86 | 3.78 | |
| pr.dry.mean | -0.02 | -0.08 | 0.03 | |
| tmax.dry.mean | -1.14 | -3.32 | 1.02 | |
| Albizia gummifera | | | | |
| intercept | 0.44 | -0.34 | 1.25 | |
| hunting1 | 6.83 | -3.67 | 17.60 | |
| logging1 | 3.33 | -8.70 | 15.80 | |
| Dist | 0.19 | -0.25 | 0.66 | |
| K | -0.02 | -0.17 | 0.12 | |
| P | 0.47 | -1.09 | 2.08 | |
| Ph | 1.75 | -5.61 | 9.05 | |
| TKN | 0.01 | -0.01 | 0.02 | |
| tmin.dry.mean | -0.91 | -3.88 | 1.99 | |

| | | | |
|---|---|---|---|
| pr.dry.mean | 0.02 | -0.04 | 0.08 |
| tmax.dry.mean | -0.31 | -2.49 | 1.88 |
| Amphimas pterocarpoides | | | |
| intercept | 0.24 | -0.48 | 0.96 |
| hunting1 | 4.88 | -4.23 | 14.20 |
| logging1 | 6.27 | -3.10 | 15.90 |
| Dist | 0.23 | -0.16 | 0.63 |
| K | 0.05 | -0.09 | 0.20 |
| P | -0.41 | -1.91 | 1.06 |
| Ph | 0.94 | -5.55 | 7.45 |
| TKN | 0.00 | -0.01 | 0.01 |
| tmin.dry.mean | 0.61 | -2.13 | 3.36 |
| pr.dry.mean | -0.03 | -0.09 | 0.03 |
| tmax.dry.mean | -0.94 | -3.12 | 1.17 |
| Angylocalyx pynaertii | | | |
| intercept | 0.16 | -0.58 | 0.89 |
| hunting1 | 2.68 | -6.53 | 11.80 |
| logging1 | 5.76 | -2.92 | 14.50 |
| Dist | 0.30 | -0.07 | 0.66 |
| K | 0.04 | -0.09 | 0.18 |
| P | 0.11 | -1.24 | 1.46 |
| Ph | 1.21 | -5.35 | 7.75 |
| TKN | -0.01 | -0.02 | 0.00 |
| tmin.dry.mean | 0.78 | -1.86 | 3.43 |
| pr.dry.mean | -0.01 | -0.07 | 0.04 |
| tmax.dry.mean | -0.97 | -2.95 | 1.01 |
| Anonidium mannii | | | |
| intercept | 0.05 | -0.65 | 0.77 |
| hunting1 | 3.05 | -5.70 | 11.70 |
| logging1 | -1.98 | -10.10 | 6.10 |
| Dist | 0.00 | -0.35 | 0.35 |
| K | -0.07 | -0.20 | 0.07 |
| P | 0.95 | -0.36 | 2.27 |
| Ph | -1.71 | -8.07 | 4.64 |
| TKN | 0.00 | -0.01 | 0.01 |
| tmin.dry.mean | -0.77 | -3.31 | 1.75 |
| pr.dry.mean | 0.02 | -0.03 | 0.07 |
| tmax.dry.mean | 0.94 | -1.00 | 2.88 |
| Anthonotha macrophylla | | | |

| | | | | |
|---|---|---|---|---|
| intercept | -0.28 | -1.22 | 0.64 | |
| hunting1 | 6.33 | -5.41 | 18.20 | |
| logging1 | 1.26 | -13.50 | 16.30 | |
| Dist | 0.02 | -0.50 | 0.56 | |
| K | 0.14 | -0.03 | 0.33 | |
| P | 0.19 | -1.64 | 2.01 | |
| Ph | -9.32 | -19.40 | 0.19 | |
| TKN | 0.00 | -0.02 | 0.01 | |
| tmin.dry.mean | -5.22 | -9.44 | -1.17 | * |
| pr.dry.mean | -0.01 | -0.10 | 0.08 | |
| tmax.dry.mean | 4.11 | 1.12 | 7.19 | * |
| Antides malaciniatum | | | | |
| intercept | 0.01 | -0.79 | 0.80 | |
| hunting1 | -3.66 | -14.00 | 6.38 | |
| logging1 | -4.76 | -13.40 | 3.72 | |
| Dist | -0.31 | -0.70 | 0.07 | |
| K | 0.07 | -0.07 | 0.22 | |
| P | -0.50 | -1.95 | 0.96 | |
| Ph | 2.25 | -4.73 | 9.18 | |
| TKN | 0.00 | -0.01 | 0.01 | |
| tmin.dry.mean | 0.60 | -2.17 | 3.37 | |
| pr.dry.mean | -0.02 | -0.08 | 0.04 | |
| tmax.dry.mean | -0.36 | -2.42 | 1.73 | |
| Barteria fistulosa | | | | |
| intercept | 0.32 | -0.52 | 1.16 | |
| hunting1 | 4.47 | -5.88 | 15.30 | |
| logging1 | -10.40 | -22.60 | 0.64 | |
| Dist | -0.25 | -0.64 | 0.13 | |
| K | 0.00 | -0.17 | 0.17 | |
| P | 1.38 | -0.34 | 3.17 | |
| Ph | 0.88 | -6.70 | 8.21 | |
| TKN | 0.01 | -0.01 | 0.02 | |
| tmin.dry.mean | 0.07 | -3.06 | 3.13 | |
| pr.dry.mean | 0.01 | -0.05 | 0.08 | |
| tmax.dry.mean | -0.34 | -2.64 | 1.98 | |
| Beilschmiedia sp. | | | | |
| intercept | 0.50 | -0.48 | 1.50 | |
| hunting1 | 2.06 | -9.71 | 13.70 | |
| logging1 | -7.17 | -17.70 | 3.04 | |

| | | | | |
|---|---|---|---|---|
| Dist | -0.15 | -0.62 | 0.31 | |
| K | 0.09 | -0.07 | 0.25 | |
| P | 0.31 | -1.31 | 1.95 | |
| Ph | 4.21 | -3.71 | 12.60 | |
| TKN | -0.01 | -0.03 | 0.00 | * |
| tmin.dry.mean | -1.23 | -4.76 | 2.36 | |
| pr.dry.mean | -0.06 | -0.13 | 0.01 | |
| tmax.dry.mean | 0.80 | -1.97 | 3.51 | |
| Blighia welwitschii | | | | |
| intercept | 0.38 | -0.37 | 1.15 | |
| hunting1 | 5.66 | -3.84 | 15.20 | |
| logging1 | 4.15 | -5.07 | 13.40 | |
| Dist | 0.41 | 0.00 | 0.83 | * |
| K | 0.11 | -0.05 | 0.28 | |
| P | -0.49 | -1.94 | 0.95 | |
| Ph | 2.38 | -4.20 | 9.03 | |
| TKN | -0.01 | -0.02 | 0.01 | |
| tmin.dry.mean | 0.35 | -2.37 | 3.09 | |
| pr.dry.mean | 0.01 | -0.04 | 0.07 | |
| tmax.dry.mean | -1.42 | -3.57 | 0.68 | |
| Camptostylus mannii | | | | |
| intercept | 0.21 | -0.53 | 0.95 | |
| hunting1 | 1.80 | -7.82 | 11.20 | |
| logging1 | 0.72 | -7.60 | 9.13 | |
| Dist | 0.09 | -0.29 | 0.48 | |
| K | -0.09 | -0.25 | 0.05 | |
| P | -0.11 | -1.51 | 1.30 | |
| Ph | 1.26 | -5.38 | 7.84 | |
| TKN | -0.01 | -0.02 | 0.00 | |
| tmin.dry.mean | -0.70 | -3.27 | 1.87 | |
| pr.dry.mean | 0.01 | -0.05 | 0.06 | |
| tmax.dry.mean | 0.82 | -1.15 | 2.81 | |
| Carapa procera | | | | |
| intercept | 0.08 | -0.68 | 0.86 | |
| hunting1 | 4.48 | -4.96 | 14.20 | |
| logging1 | 1.89 | -6.71 | 10.80 | |
| Dist | 0.01 | -0.37 | 0.40 | |
| K | 0.09 | -0.05 | 0.25 | |
| P | 0.45 | -0.96 | 1.88 | |

| | | | | |
|---|---|---|---|---|
| Ph | -0.84 | -7.78 | 6.13 | |
| TKN | 0.00 | -0.01 | 0.01 | |
| tmin.dry.mean | 1.81 | -0.92 | 4.54 | |
| pr.dry.mean | -0.02 | -0.08 | 0.04 | |
| tmax.dry.mean | -1.29 | -3.39 | 0.80 | |
| Celtis adolfi friderici | | | | |
| intercept | 0.34 | -0.39 | 1.06 | |
| hunting1 | -1.87 | -10.80 | 7.05 | |
| logging1 | -8.20 | -16.50 | 0.00 | |
| Dist | -0.47 | -0.83 | -0.12 | * |
| K | -0.18 | -0.33 | -0.03 | * |
| P | 0.28 | -1.06 | 1.61 | |
| Ph | 4.67 | -1.84 | 11.20 | |
| TKN | 0.00 | -0.01 | 0.01 | |
| tmin.dry.mean | 0.31 | -2.34 | 2.93 | |
| pr.dry.mean | -0.01 | -0.07 | 0.04 | |
| tmax.dry.mean | 0.71 | -1.30 | 2.77 | |
| Celtis mildbraedii | | | | |
| intercept | 1.46 | 0.18 | 2.80 | * |
| hunting1 | 11.40 | -3.01 | 26.30 | |
| logging1 | 9.80 | -3.03 | 23.90 | |
| Dist | 0.88 | 0.30 | 1.50 | * |
| K | 0.17 | -0.14 | 0.48 | |
| P | 2.20 | -0.60 | 5.13 | |
| Ph | 11.40 | -2.18 | 25.10 | |
| TKN | -0.05 | -0.07 | -0.02 | * |
| tmin.dry.mean | -4.63 | -10.30 | 1.00 | |
| pr.dry.mean | 0.01 | -0.10 | 0.12 | |
| tmax.dry.mean | 0.84 | -3.47 | 5.14 | |
| Chrysophyllum boukokoense | | | | |
| intercept | 0.02 | -0.77 | 0.81 | |
| hunting1 | 6.78 | -2.94 | 16.80 | |
| logging1 | 2.51 | -7.16 | 12.60 | |
| Dist | 0.28 | -0.13 | 0.71 | |
| K | -0.04 | -0.19 | 0.11 | |
| P | 0.12 | -1.37 | 1.64 | |
| Ph | -4.11 | -11.90 | 3.41 | |
| TKN | -0.01 | -0.03 | 0.00 | |
| tmin.dry.mean | -0.99 | -3.86 | 1.85 | |

| | | | |
|---|---|---|---|
| pr.dry.mean | 0.00 | -0.06 | 0.06 |
| tmax.dry.mean | 1.21 | -1.05 | 3.51 |
| Chrysophyllum lacourtiana | | | |
| intercept | 0.75 | -0.12 | 1.66 |
| hunting1 | 3.79 | -6.47 | 14.00 |
| logging1 | -8.64 | -19.90 | 1.85 |
| Dist | -0.35 | -0.78 | 0.07 |
| K | -0.14 | -0.30 | 0.02 |
| P | 1.23 | -0.37 | 2.86 |
| Ph | 5.97 | -1.31 | 13.50 |
| TKN | 0.00 | -0.02 | 0.01 |
| tmin.dry.mean | -1.83 | -6.30 | 2.04 |
| pr.dry.mean | -0.04 | -0.12 | 0.04 |
| tmax.dry.mean | 1.52 | -1.34 | 4.85 |
| Cleistopholis patens | | | |
| intercept | -0.35 | -1.10 | 0.38 |
| hunting1 | 3.22 | -6.35 | 12.90 |
| logging1 | 6.04 | -3.06 | 15.40 |
| Dist | 0.25 | -0.13 | 0.65 |
| K | -0.02 | -0.17 | 0.13 |
| P | 0.71 | -0.77 | 2.23 |
| Ph | -5.85 | -13.00 | 1.13 |
| TKN | -0.01 | -0.03 | 0.00 |
| tmin.dry.mean | 1.48 | -1.20 | 4.22 |
| pr.dry.mean | -0.04 | -0.09 | 0.02 |
| tmax.dry.mean | -0.17 | -2.27 | 1.95 |
| Cola lateritia | | | |
| intercept | 0.53 | -0.19 | 1.25 |
| hunting1 | 5.79 | -2.95 | 14.70 |
| logging1 | 2.52 | -5.66 | 10.70 |
| Dist | 0.15 | -0.20 | 0.51 |
| K | -0.02 | -0.15 | 0.12 |
| P | -0.02 | -1.33 | 1.31 |
| Ph | 3.48 | -2.88 | 9.89 |
| TKN | 0.00 | -0.02 | 0.01 |
| tmin.dry.mean | -0.27 | -2.93 | 2.34 |
| pr.dry.mean | -0.02 | -0.08 | 0.03 |
| tmax.dry.mean | -0.08 | -2.08 | 1.95 |
| Dacryodes edulis | | | |

| | | | | |
|---|---|---|---|---|
| intercept | -0.26 | -1.04 | 0.51 | |
| hunting1 | -4.73 | -14.90 | 5.12 | |
| logging1 | -2.80 | -11.90 | 6.07 | |
| Dist | -0.21 | -0.63 | 0.19 | |
| K | 0.05 | -0.10 | 0.21 | |
| P | -0.66 | -2.14 | 0.79 | |
| Ph | -0.63 | -7.37 | 6.14 | |
| TKN | -0.01 | -0.02 | 0.01 | |
| tmin.dry.mean | 0.42 | -2.33 | 3.10 | |
| pr.dry.mean | 0.02 | -0.04 | 0.08 | |
| tmax.dry.mean | 0.06 | -2.01 | 2.17 | |
| Desplatsia chrysochlamys | | | | |
| intercept | 0.20 | -0.97 | 1.38 | |
| hunting1 | -5.72 | -20.40 | 7.87 | |
| logging1 | -14.40 | -26.90 | -3.02 | * |
| Dist | -0.73 | -1.29 | -0.23 | * |
| K | 0.02 | -0.15 | 0.19 | |
| P | 0.01 | -1.86 | 1.88 | |
| Ph | 3.78 | -4.85 | 12.90 | |
| TKN | -0.01 | -0.02 | 0.01 | |
| tmin.dry.mean | -2.81 | -7.46 | 1.67 | |
| pr.dry.mean | -0.03 | -0.11 | 0.06 | |
| tmax.dry.mean | 2.70 | -0.61 | 6.07 | |
| Desplatsia dewevrei | | | | |
| intercept | 0.65 | -0.40 | 1.78 | |
| hunting1 | 0.16 | -12.70 | 13.00 | |
| logging1 | -10.30 | -23.20 | 1.27 | |
| Dist | -0.45 | -0.94 | 0.01 | |
| K | 0.04 | -0.11 | 0.19 | |
| P | -0.02 | -1.77 | 1.79 | |
| Ph | 7.09 | -0.94 | 15.70 | |
| TKN | 0.00 | -0.01 | 0.01 | |
| tmin.dry.mean | -1.75 | -5.90 | 2.26 | |
| pr.dry.mean | -0.03 | -0.10 | 0.05 | |
| tmax.dry.mean | 0.88 | -2.21 | 4.01 | |
| Dialium pachyphyllum | | | | |
| intercept | 0.40 | -0.30 | 1.10 | |
| hunting1 | 7.70 | -0.97 | 16.50 | |
| logging1 | -3.11 | -11.20 | 5.02 | |

| | | | | |
|---|---|---|---|---|
| Dist | 0.29 | -0.06 | 0.63 | |
| K | -0.08 | -0.22 | 0.06 | |
| P | 0.58 | -0.75 | 1.91 | |
| Ph | -0.44 | -6.79 | 5.87 | |
| TKN | -0.01 | -0.02 | 0.01 | |
| tmin.dry.mean | -1.58 | -4.10 | 0.95 | |
| pr.dry.mean | 0.00 | -0.05 | 0.05 | |
| tmax.dry.mean | 1.42 | -0.51 | 3.36 | |
| Dichostemma glaucescens | | | | |
| intercept | -4.18 | -6.31 | -2.25 | * |
| hunting1 | 25.50 | 9.75 | 38.40 | * |
| logging1 | 28.80 | 11.90 | 34.10 | * |
| Dist | 1.81 | 1.19 | 2.21 | * |
| K | 1.40 | 0.87 | 1.92 | * |
| P | -5.73 | -10.50 | -1.46 | * |
| Ph | -71.70 | -97.50 | -47.50 | * |
| TKN | -0.06 | -0.10 | -0.03 | * |
| tmin.dry.mean | -11.60 | -21.30 | -2.45 | * |
| pr.dry.mean | 0.11 | -0.06 | 0.30 | |
| tmax.dry.mean | 11.40 | 4.30 | 19.00 | * |
| Diospyros bipindensis | | | | |
| intercept | 0.04 | -0.68 | 0.76 | |
| hunting1 | -0.07 | -9.08 | 8.98 | |
| logging1 | -2.67 | -10.90 | 5.54 | |
| Dist | 0.38 | 0.02 | 0.74 | * |
| K | 0.22 | 0.08 | 0.36 | * |
| P | 0.20 | -1.14 | 1.54 | |
| Ph | -0.12 | -6.68 | 6.42 | |
| TKN | -0.02 | -0.03 | 0.00 | * |
| tmin.dry.mean | -1.51 | -4.12 | 1.09 | |
| pr.dry.mean | -0.02 | -0.07 | 0.03 | |
| tmax.dry.mean | 0.70 | -1.31 | 2.68 | |
| Diospyros canaliculata | | | | |
| intercept | 0.70 | -0.24 | 1.65 | |
| hunting1 | -5.27 | -16.50 | 5.96 | |
| logging1 | 2.02 | -7.47 | 12.10 | |
| Dist | -0.04 | -0.46 | 0.40 | |
| K | -0.14 | -0.35 | 0.06 | |
| P | 1.02 | -0.93 | 2.99 | |

| | | | | |
|---|---|---|---|---|
| Ph | 12.60 | 3.43 | 22.00 | * |
| TKN | 0.01 | -0.01 | 0.02 | |
| tmin.dry.mean | 0.68 | -3.20 | 4.53 | |
| pr.dry.mean | -0.05 | -0.13 | 0.02 | |
| tmax.dry.mean | -1.38 | -4.33 | 1.55 | |
| Diospyros crassiflora | | | | |
| intercept | 0.08 | -0.77 | 0.92 | |
| hunting1 | 5.57 | -4.65 | 15.90 | |
| logging1 | -10.60 | -22.50 | 0.37 | |
| Dist | 0.00 | -0.40 | 0.40 | |
| K | 0.05 | -0.11 | 0.22 | |
| P | 1.04 | -0.55 | 2.68 | |
| Ph | -3.91 | -11.90 | 3.75 | |
| TKN | -0.02 | -0.03 | 0.00 | * |
| tmin.dry.mean | -2.17 | -5.40 | 0.95 | |
| pr.dry.mean | 0.03 | -0.03 | 0.10 | |
| tmax.dry.mean | 1.87 | -0.44 | 4.30 | |
| Diospyros iturensis | | | | |
| intercept | 0.41 | -0.31 | 1.15 | |
| hunting1 | 5.53 | -3.60 | 14.80 | |
| logging1 | 2.15 | -6.00 | 10.30 | |
| Dist | 0.39 | 0.03 | 0.76 | * |
| K | -0.09 | -0.23 | 0.05 | |
| P | 1.11 | -0.25 | 2.47 | |
| Ph | 2.22 | -4.30 | 8.80 | |
| TKN | 0.00 | -0.02 | 0.01 | |
| tmin.dry.mean | 0.28 | -2.29 | 2.84 | |
| pr.dry.mean | -0.05 | -0.10 | 0.00 | |
| tmax.dry.mean | -0.35 | -2.32 | 1.61 | |
| Diospyros mannii | | | | |
| intercept | -0.04 | -0.83 | 0.74 | |
| hunting1 | -3.59 | -14.40 | 6.88 | |
| logging1 | 1.44 | -7.12 | 10.10 | |
| Dist | 0.05 | -0.37 | 0.47 | |
| K | -0.01 | -0.15 | 0.14 | |
| P | -0.72 | -2.29 | 0.81 | |
| Ph | 1.98 | -4.70 | 8.79 | |
| TKN | -0.01 | -0.02 | 0.01 | |
| tmin.dry.mean | 0.65 | -2.10 | 3.39 | |

| | | | | |
|---|---|---|---|---|
| pr.dry.mean | 0.03 | -0.03 | 0.09 | |
| tmax.dry.mean | -0.61 | -2.69 | 1.48 | |
| Discoglypremna caloneura | | | | |
| intercept | -0.32 | -1.06 | 0.42 | |
| hunting1 | -2.90 | -12.10 | 6.39 | |
| logging1 | 2.41 | -5.81 | 10.70 | |
| Dist | -0.10 | -0.47 | 0.27 | |
| K | -0.02 | -0.17 | 0.12 | |
| P | -0.26 | -1.58 | 1.09 | |
| Ph | -2.38 | -8.86 | 4.08 | |
| TKN | 0.00 | -0.01 | 0.01 | |
| tmin.dry.mean | 0.20 | -2.56 | 2.95 | |
| pr.dry.mean | -0.03 | -0.08 | 0.03 | |
| tmax.dry.mean | 0.66 | -1.39 | 2.73 | |
| Drypetes gossweileri | | | | |
| intercept | 0.32 | -0.43 | 1.07 | |
| hunting1 | 4.39 | -5.57 | 14.30 | |
| logging1 | 0.87 | -8.34 | 10.10 | |
| Dist | 0.32 | -0.09 | 0.73 | |
| K | -0.05 | -0.20 | 0.10 | |
| P | 0.40 | -1.13 | 1.96 | |
| Ph | 1.78 | -5.10 | 8.68 | |
| TKN | -0.01 | -0.03 | 0.01 | |
| tmin.dry.mean | 0.23 | -2.67 | 3.17 | |
| pr.dry.mean | 0.03 | -0.04 | 0.09 | |
| tmax.dry.mean | -0.61 | -2.87 | 1.59 | |
| Drypetes ituriensis | | | | |
| intercept | -0.02 | -0.98 | 0.92 | |
| hunting1 | -1.22 | -13.60 | 10.40 | |
| logging1 | 9.70 | -0.21 | 20.20 | |
| Dist | 0.53 | 0.07 | 1.01 | * |
| K | 0.08 | -0.09 | 0.25 | |
| P | -1.44 | -3.36 | 0.31 | |
| Ph | 1.29 | -6.63 | 9.39 | |
| TKN | -0.01 | -0.03 | 0.01 | |
| tmin.dry.mean | -0.13 | -3.11 | 2.79 | |
| pr.dry.mean | 0.00 | -0.07 | 0.06 | |
| tmax.dry.mean | -0.47 | -2.71 | 1.79 | |
| Drypetes occidentalis | | | | |

| | | | | |
|---|---|---|---|---|
| intercept | 0.57 | -0.20 | 1.37 | |
| hunting1 | 7.07 | -1.88 | 16.20 | |
| logging1 | 3.06 | -5.60 | 11.70 | |
| Dist | 0.36 | -0.02 | 0.75 | |
| K | 0.04 | -0.10 | 0.19 | |
| P | -0.44 | -1.83 | 0.96 | |
| Ph | 2.62 | -4.41 | 9.69 | |
| TKN | 0.00 | -0.02 | 0.01 | |
| tmin.dry.mean | -2.48 | -5.69 | 0.45 | |
| pr.dry.mean | -0.04 | -0.10 | 0.02 | |
| tmax.dry.mean | 1.18 | -1.08 | 3.65 | |
| Drypetes polyantha | | | | |
| intercept | 0.42 | -0.33 | 1.18 | |
| hunting1 | -0.55 | -10.00 | 8.87 | |
| logging1 | -0.43 | -8.78 | 7.97 | |
| Dist | 0.10 | -0.27 | 0.47 | |
| K | -0.06 | -0.22 | 0.09 | |
| P | -0.30 | -1.75 | 1.15 | |
| Ph | 5.14 | -1.78 | 12.30 | |
| TKN | 0.00 | -0.01 | 0.01 | |
| tmin.dry.mean | -1.72 | -4.62 | 1.12 | |
| pr.dry.mean | 0.00 | -0.06 | 0.06 | |
| tmax.dry.mean | 0.76 | -1.36 | 2.93 | |
| Drypetes sp. | | | | |
| intercept | 0.15 | -0.64 | 0.94 | |
| hunting1 | 11.60 | 1.13 | 22.40 | * |
| logging1 | 12.00 | 0.19 | 25.30 | * |
| Dist | 0.67 | 0.20 | 1.19 | * |
| K | -0.17 | -0.37 | 0.01 | |
| P | 0.80 | -0.92 | 2.56 | |
| Ph | -4.01 | -12.30 | 3.91 | |
| TKN | 0.00 | -0.02 | 0.02 | |
| tmin.dry.mean | 0.63 | -2.54 | 3.80 | |
| pr.dry.mean | 0.01 | -0.05 | 0.07 | |
| tmax.dry.mean | -0.54 | -2.84 | 1.75 | |
| Duboscia macrocarpa | | | | |
| intercept | 0.25 | -0.53 | 1.05 | |
| hunting1 | 3.31 | -6.76 | 13.70 | |
| logging1 | 0.41 | -8.72 | 9.68 | |

| | | | |
|---|---|---|---|
| Dist | 0.07 | -0.31 | 0.47 |
| K | -0.11 | -0.26 | 0.04 |
| P | 0.87 | -0.69 | 2.49 |
| Ph | 1.48 | -5.21 | 8.34 |
| TKN | 0.01 | 0.00 | 0.02 |
| tmin.dry.mean | 0.48 | -2.27 | 3.29 |
| pr.dry.mean | 0.00 | -0.06 | 0.06 |
| tmax.dry.mean | -0.65 | -2.81 | 1.46 |
| Entandrophragma angolense | | | |
| intercept | -0.34 | -1.16 | 0.47 |
| hunting1 | 1.45 | -9.10 | 11.80 |
| logging1 | 4.83 | -5.07 | 15.20 |
| Dist | 0.08 | -0.36 | 0.52 |
| K | 0.12 | -0.04 | 0.27 |
| P | -0.02 | -1.49 | 1.44 |
| Ph | -4.54 | -12.30 | 2.92 |
| TKN | 0.00 | -0.01 | 0.01 |
| tmin.dry.mean | 0.95 | -1.92 | 3.85 |
| pr.dry.mean | -0.02 | -0.09 | 0.04 |
| tmax.dry.mean | -0.49 | -2.66 | 1.67 |
| Entandrophragma candollei | | | |
| intercept | 0.17 | -0.60 | 0.96 |
| hunting1 | 2.49 | -7.30 | 12.20 |
| logging1 | 3.45 | -5.03 | 12.10 |
| Dist | 0.10 | -0.28 | 0.48 |
| K | 0.03 | -0.11 | 0.17 |
| P | -0.11 | -1.50 | 1.29 |
| Ph | 1.45 | -5.35 | 8.53 |
| TKN | 0.00 | -0.02 | 0.01 |
| tmin.dry.mean | 1.43 | -1.19 | 4.07 |
| pr.dry.mean | -0.02 | -0.08 | 0.03 |
| tmax.dry.mean | -1.13 | -3.17 | 0.88 |
| Entandrophragma cylindricum | | | |
| intercept | 0.27 | -0.47 | 1.01 |
| hunting1 | 0.26 | -8.70 | 9.12 |
| logging1 | -0.79 | -9.08 | 7.51 |
| Dist | 0.13 | -0.24 | 0.48 |
| K | -0.04 | -0.19 | 0.10 |
| P | 0.00 | -1.36 | 1.39 |

| | | | | |
|---|---|---|---|---|
| Ph | 3.09 | -3.65 | 9.93 | |
| TKN | 0.00 | -0.02 | 0.01 | |
| tmin.dry.mean | -0.56 | -3.13 | 2.02 | |
| pr.dry.mean | 0.01 | -0.04 | 0.06 | |
| tmax.dry.mean | 0.18 | -1.81 | 2.18 | |
| Entandrophragma utile | | | | |
| intercept | 0.29 | -0.52 | 1.09 | |
| hunting1 | -0.45 | -10.70 | 9.55 | |
| logging1 | -2.90 | -12.50 | 6.67 | |
| Dist | -0.06 | -0.49 | 0.36 | |
| K | -0.01 | -0.16 | 0.14 | |
| P | 0.32 | -1.19 | 1.81 | |
| Ph | 3.45 | -3.63 | 10.70 | |
| TKN | -0.01 | -0.02 | 0.00 | |
| tmin.dry.mean | -0.66 | -3.70 | 2.33 | |
| pr.dry.mean | -0.03 | -0.10 | 0.03 | |
| tmax.dry.mean | 0.45 | -1.91 | 2.82 | |
| Erythrophleum suaveolens | | | | |
| intercept | 0.84 | 0.03 | 1.67 | * |
| hunting1 | 9.05 | -0.23 | 18.60 | |
| logging1 | -1.66 | -10.40 | 6.96 | |
| Dist | 0.21 | -0.16 | 0.58 | |
| K | -0.05 | -0.20 | 0.10 | |
| P | -0.30 | -1.66 | 1.05 | |
| Ph | 5.40 | -1.73 | 12.80 | |
| TKN | 0.00 | -0.01 | 0.01 | |
| tmin.dry.mean | -0.28 | -3.00 | 2.35 | |
| pr.dry.mean | 0.00 | -0.06 | 0.05 | |
| tmax.dry.mean | -0.43 | -2.48 | 1.63 | |
| Fernandoa adolfi friderici | | | | |
| intercept | 0.63 | -0.28 | 1.55 | |
| hunting1 | 5.61 | -6.25 | 17.40 | |
| logging1 | 0.99 | -9.07 | 11.30 | |
| Dist | 0.15 | -0.29 | 0.60 | |
| K | -0.11 | -0.27 | 0.05 | |
| P | 0.15 | -1.58 | 1.94 | |
| Ph | 4.76 | -3.12 | 12.80 | |
| TKN | 0.00 | -0.01 | 0.02 | |
| tmin.dry.mean | -0.62 | -3.63 | 2.37 | |

| | | | | |
|---|---|---|---|---|
| pr.dry.mean | 0.01 | -0.06 | 0.07 | |
| tmax.dry.mean | -0.13 | -2.51 | 2.19 | |
| Funtumia elastica | | | | |
| intercept | -0.01 | -0.72 | 0.70 | |
| hunting1 | -5.40 | -14.20 | 3.23 | |
| logging1 | -5.11 | -13.60 | 3.16 | |
| Dist | -0.26 | -0.63 | 0.08 | |
| K | -0.03 | -0.17 | 0.10 | |
| P | -0.04 | -1.35 | 1.27 | |
| Ph | 2.58 | -3.87 | 9.06 | |
| TKN | 0.00 | -0.01 | 0.01 | |
| tmin.dry.mean | 0.16 | -2.40 | 2.71 | |
| pr.dry.mean | -0.02 | -0.07 | 0.03 | |
| tmax.dry.mean | 0.39 | -1.57 | 2.35 | |
| Garcinia punctata | | | | |
| intercept | 0.15 | -0.80 | 1.03 | |
| hunting1 | 17.90 | 6.78 | 27.70 | * |
| logging1 | 14.30 | -0.01 | 28.50 | |
| Dist | 1.19 | 0.57 | 1.78 | * |
| K | 0.27 | 0.05 | 0.51 | * |
| P | -2.19 | -4.32 | -0.16 | * |
| Ph | -7.82 | -19.50 | 2.48 | |
| TKN | -0.02 | -0.05 | 0.00 | * |
| tmin.dry.mean | -1.47 | -5.42 | 2.20 | |
| pr.dry.mean | 0.06 | -0.01 | 0.15 | |
| tmax.dry.mean | -0.18 | -3.20 | 2.94 | |
| Greenwayodendron suaveolens | | | | |
| intercept | 1.81 | 0.95 | 2.63 | * |
| hunting1 | 21.50 | 11.00 | 28.20 | * |
| logging1 | -0.51 | -11.50 | 11.10 | |
| Dist | 0.86 | 0.41 | 1.31 | * |
| K | -0.26 | -0.46 | -0.08 | * |
| P | 2.78 | 0.99 | 4.46 | * |
| Ph | 10.60 | 2.14 | 19.40 | * |
| TKN | -0.01 | -0.03 | 0.00 | |
| tmin.dry.mean | -0.53 | -4.01 | 2.82 | |
| pr.dry.mean | 0.07 | 0.00 | 0.14 | |
| tmax.dry.mean | -1.42 | -3.93 | 1.16 | |
| Grossera macrantha | | | | |

| | | | | |
|---|---|---|---|---|
| intercept | -0.01 | -2.09 | 2.17 | |
| hunting1 | -9.17 | -23.90 | 10.30 | |
| logging1 | -27.90 | -35.60 | -9.29 | * |
| Dist | 0.09 | -0.59 | 1.02 | |
| K | 0.47 | -0.09 | 1.09 | |
| P | -0.81 | -5.49 | 3.96 | |
| Ph | 1.76 | -24.30 | 26.80 | |
| TKN | -0.05 | -0.09 | -0.01 | * |
| tmin.dry.mean | -4.70 | -14.60 | 5.07 | |
| pr.dry.mean | -0.07 | -0.28 | 0.13 | |
| tmax.dry.mean | 3.53 | -4.03 | 11.00 | |
| Guarea cedrata | | | | |
| intercept | 0.41 | -0.35 | 1.17 | |
| hunting1 | 5.67 | -4.16 | 15.60 | |
| logging1 | -3.81 | -12.70 | 4.86 | |
| Dist | 0.11 | -0.27 | 0.49 | |
| K | -0.09 | -0.27 | 0.07 | |
| P | 0.31 | -1.19 | 1.86 | |
| Ph | 1.13 | -5.75 | 8.03 | |
| TKN | 0.00 | -0.02 | 0.01 | |
| tmin.dry.mean | -1.20 | -3.96 | 1.56 | |
| pr.dry.mean | 0.01 | -0.05 | 0.06 | |
| tmax.dry.mean | 0.91 | -1.22 | 3.03 | |
| Guarea thompsonii | | | | |
| intercept | -0.49 | -1.22 | 0.24 | |
| hunting1 | 9.19 | -0.17 | 18.60 | |
| logging1 | 10.30 | 1.20 | 19.70 | * |
| Dist | 0.58 | 0.21 | 0.98 | * |
| K | 0.10 | -0.04 | 0.24 | |
| P | 0.02 | -1.39 | 1.46 | |
| Ph | -12.00 | -19.10 | -5.08 | * |
| TKN | -0.01 | -0.03 | 0.00 | |
| tmin.dry.mean | -1.86 | -4.54 | 0.81 | |
| pr.dry.mean | 0.00 | -0.05 | 0.06 | |
| tmax.dry.mean | 2.06 | 0.01 | 4.09 | * |
| Hannoa klaineana | | | | |
| intercept | -0.25 | -1.07 | 0.56 | |
| hunting1 | -2.12 | -11.60 | 7.24 | |
| logging1 | -1.37 | -11.00 | 8.20 | |

| | | | | |
|---|---|---|---|---|
| Dist | -0.02 | -0.43 | 0.39 | |
| K | 0.00 | -0.18 | 0.16 | |
| P | -0.25 | -1.68 | 1.17 | |
| Ph | -2.88 | -10.80 | 4.84 | |
| TKN | 0.00 | -0.01 | 0.01 | |
| tmin.dry.mean | -2.08 | -5.01 | 0.80 | |
| pr.dry.mean | 0.03 | -0.03 | 0.09 | |
| tmax.dry.mean | 1.72 | -0.45 | 3.95 | |
| Hexalobus crispiflorus | | | | |
| intercept | -0.34 | -1.14 | 0.44 | |
| hunting1 | -4.97 | -15.00 | 4.67 | |
| logging1 | -0.59 | -8.83 | 7.67 | |
| Dist | -0.16 | -0.55 | 0.22 | |
| K | -0.02 | -0.16 | 0.13 | |
| P | -0.19 | -1.55 | 1.16 | |
| Ph | -1.39 | -8.33 | 5.40 | |
| TKN | 0.00 | -0.01 | 0.01 | |
| tmin.dry.mean | 0.53 | -2.06 | 3.10 | |
| pr.dry.mean | 0.03 | -0.03 | 0.08 | |
| tmax.dry.mean | 0.06 | -1.96 | 2.07 | |
| Irvingia grandifolia | | | | |
| intercept | 0.57 | -0.38 | 1.57 | |
| hunting1 | 4.97 | -5.83 | 16.00 | |
| logging1 | -6.07 | -16.00 | 3.62 | |
| Dist | -0.01 | -0.43 | 0.40 | |
| K | 0.01 | -0.18 | 0.19 | |
| P | 0.90 | -0.79 | 2.64 | |
| Ph | 4.50 | -4.39 | 13.50 | |
| TKN | 0.00 | -0.01 | 0.01 | |
| tmin.dry.mean | 0.94 | -2.03 | 4.00 | |
| pr.dry.mean | -0.03 | -0.09 | 0.03 | |
| tmax.dry.mean | -1.31 | -3.83 | 1.10 | |
| Isolona hexaloba | | | | |
| intercept | 0.57 | -0.26 | 1.44 | |
| hunting1 | 11.70 | 1.97 | 21.80 | * |
| logging1 | 5.06 | -4.14 | 14.50 | |
| Dist | 0.40 | 0.02 | 0.79 | * |
| K | -0.02 | -0.17 | 0.12 | |
| P | 0.04 | -1.39 | 1.50 | |

| | | | |
|---|---|---|---|
| Ph | 1.50 | -6.32 | 9.46 |
| TKN | 0.00 | -0.02 | 0.01 |
| tmin.dry.mean | 1.70 | -1.20 | 4.66 |
| pr.dry.mean | -0.03 | -0.09 | 0.03 |
| tmax.dry.mean | -1.62 | -4.09 | 0.72 |
| Keayodendron bridelioides | | | |
| intercept | 0.32 | -0.56 | 1.21 |
| hunting1 | 5.33 | -7.71 | 17.80 |
| logging1 | 9.95 | -2.16 | 24.00 |
| Dist | 0.13 | -0.43 | 0.68 |
| K | -0.10 | -0.25 | 0.05 |
| P | 0.50 | -1.13 | 2.15 |
| Ph | 1.35 | -6.39 | 9.00 |
| TKN | 0.00 | -0.01 | 0.02 |
| tmin.dry.mean | -0.67 | -4.97 | 3.52 |
| pr.dry.mean | -0.01 | -0.09 | 0.07 |
| tmax.dry.mean | 0.06 | -2.99 | 3.24 |
| Klainedoxa gabonensis | | | |
| intercept | -0.49 | -1.34 | 0.33 |
| hunting1 | -2.41 | -13.30 | 7.97 |
| logging1 | 1.54 | -7.49 | 10.70 |
| Dist | -0.17 | -0.60 | 0.26 |
| K | 0.01 | -0.16 | 0.18 |
| P | -0.53 | -2.10 | 1.00 |
| Ph | -4.84 | -12.60 | 2.62 |
| TKN | 0.00 | -0.01 | 0.01 |
| tmin.dry.mean | 0.47 | -2.42 | 3.34 |
| pr.dry.mean | -0.01 | -0.07 | 0.05 |
| tmax.dry.mean | 0.60 | -1.61 | 2.84 |
| Lecaniodiscus cupanioides | | | |
| intercept | 0.36 | -0.38 | 1.11 |
| hunting1 | 4.17 | -5.31 | 13.80 |
| logging1 | 0.64 | -8.29 | 9.71 |
| Dist | 0.14 | -0.24 | 0.51 |
| K | -0.03 | -0.17 | 0.11 |
| P | 0.05 | -1.41 | 1.54 |
| Ph | 2.30 | -4.28 | 8.98 |
| TKN | 0.00 | -0.01 | 0.01 |
| tmin.dry.mean | -0.08 | -2.75 | 2.54 |

| | | | | |
|---|---|---|---|---|
| pr.dry.mean | 0.01 | -0.04 | 0.07 | |
| tmax.dry.mean | -0.40 | -2.41 | 1.64 | |
| Lepidobotrys staudtii | | | | |
| intercept | 0.13 | -0.84 | 1.14 | |
| hunting1 | 0.40 | -13.30 | 13.60 | |
| logging1 | -11.10 | -23.40 | 0.03 | |
| Dist | -0.41 | -0.94 | 0.06 | |
| K | 0.05 | -0.10 | 0.21 | |
| P | 0.38 | -1.41 | 2.27 | |
| Ph | 0.92 | -6.74 | 9.00 | |
| TKN | -0.01 | -0.02 | 0.00 | |
| tmin.dry.mean | 0.50 | -2.82 | 4.06 | |
| pr.dry.mean | 0.07 | 0.00 | 0.15 | |
| tmax.dry.mean | -0.40 | -2.99 | 2.02 | |
| Lindackeria dentata | | | | |
| intercept | 0.68 | -0.15 | 1.51 | |
| hunting1 | 8.44 | -1.57 | 18.50 | |
| logging1 | -1.31 | -10.80 | 8.08 | |
| Dist | 0.10 | -0.29 | 0.49 | |
| K | -0.05 | -0.21 | 0.12 | |
| P | 0.66 | -0.91 | 2.23 | |
| Ph | 4.54 | -2.92 | 12.10 | |
| TKN | -0.01 | -0.02 | 0.01 | |
| tmin.dry.mean | 2.61 | -0.48 | 5.77 | |
| pr.dry.mean | -0.06 | -0.13 | 0.00 | * |
| tmax.dry.mean | -1.96 | -4.48 | 0.52 | |
| Lovoa trichilioides | | | | |
| intercept | -0.12 | -0.87 | 0.65 | |
| hunting1 | 1.24 | -8.40 | 10.80 | |
| logging1 | 1.12 | -8.06 | 10.30 | |
| Dist | 0.16 | -0.22 | 0.54 | |
| K | 0.04 | -0.11 | 0.18 | |
| P | -0.38 | -1.88 | 1.13 | |
| Ph | -2.75 | -9.76 | 4.17 | |
| TKN | 0.00 | -0.01 | 0.01 | |
| tmin.dry.mean | -1.72 | -4.56 | 1.05 | |
| pr.dry.mean | 0.04 | -0.02 | 0.09 | |
| tmax.dry.mean | 1.11 | -0.95 | 3.21 | |
| Macaranga barteri | | | | |

| | | | | |
|---|---|---|---|---|
| intercept | -0.31 | -1.04 | 0.42 | |
| hunting1 | -0.74 | -9.64 | 8.24 | |
| logging1 | 1.46 | -6.97 | 9.91 | |
| Dist | 0.09 | -0.27 | 0.45 | |
| K | -0.01 | -0.15 | 0.13 | |
| P | -0.33 | -1.68 | 1.03 | |
| Ph | -3.54 | -10.30 | 3.13 | |
| TKN | -0.01 | -0.02 | 0.00 | |
| tmin.dry.mean | 0.43 | -2.15 | 3.01 | |
| pr.dry.mean | 0.00 | -0.06 | 0.05 | |
| tmax.dry.mean | 0.52 | -1.47 | 2.51 | |
| Macaranga spinosa | | | | |
| intercept | -0.19 | -1.55 | 0.94 | |
| hunting1 | 28.10 | 14.20 | 41.10 | * |
| logging1 | 20.90 | 8.57 | 31.40 | * |
| Dist | 1.48 | 0.86 | 2.03 | * |
| K | -0.04 | -0.31 | 0.26 | |
| P | 1.59 | -0.73 | 4.03 | |
| Ph | -18.60 | -35.20 | -5.58 | * |
| TKN | -0.03 | -0.06 | -0.01 | * |
| tmin.dry.mean | -1.25 | -5.20 | 2.64 | |
| pr.dry.mean | -0.03 | -0.12 | 0.05 | |
| tmax.dry.mean | 1.77 | -1.35 | 5.18 | |
| Massularia acuminata | | | | |
| intercept | -0.08 | -0.97 | 0.83 | |
| hunting1 | 0.15 | -11.60 | 11.60 | |
| logging1 | 3.72 | -6.22 | 14.10 | |
| Dist | 0.26 | -0.18 | 0.71 | |
| K | -0.02 | -0.19 | 0.14 | |
| P | -0.15 | -1.83 | 1.53 | |
| Ph | -0.53 | -8.48 | 7.70 | |
| TKN | 0.00 | -0.01 | 0.02 | |
| tmin.dry.mean | 0.31 | -2.70 | 3.33 | |
| pr.dry.mean | 0.05 | -0.02 | 0.11 | |
| tmax.dry.mean | -0.79 | -3.16 | 1.55 | |
| Monodora tenuifolia | | | | |
| intercept | 0.28 | -0.54 | 1.11 | |
| hunting1 | 7.72 | -2.01 | 17.70 | |
| logging1 | -0.55 | -10.00 | 9.02 | |

| | | | | |
|---|---|---|---|---|
| Dist | 0.25 | -0.14 | 0.64 | |
| K | -0.04 | -0.18 | 0.10 | |
| P | 0.89 | -0.57 | 2.41 | |
| Ph | -1.37 | -8.73 | 6.11 | |
| TKN | -0.01 | -0.02 | 0.01 | |
| tmin.dry.mean | -0.90 | -3.60 | 1.80 | |
| pr.dry.mean | 0.01 | -0.04 | 0.07 | |
| tmax.dry.mean | 0.47 | -1.62 | 2.57 | |
| Myrianthus arboreus | | | | |
| intercept | 0.17 | -0.91 | 1.25 | |
| hunting1 | 1.59 | -10.40 | 13.00 | |
| logging1 | -1.04 | -11.50 | 9.76 | |
| Dist | -0.66 | -1.13 | -0.20 | * |
| K | 0.09 | -0.06 | 0.24 | |
| P | 0.49 | -1.16 | 2.15 | |
| Ph | -1.41 | -9.90 | 7.10 | |
| TKN | 0.00 | -0.02 | 0.01 | |
| tmin.dry.mean | -5.62 | -11.40 | -0.13 | * |
| pr.dry.mean | -0.13 | -0.25 | -0.03 | * |
| tmax.dry.mean | 5.17 | 1.13 | 9.39 | * |
| Nesogordonia kabingaensis | | | | |
| intercept | -0.43 | -1.15 | 0.28 | |
| hunting1 | 0.57 | -8.20 | 9.48 | |
| logging1 | 4.08 | -4.10 | 12.30 | |
| Dist | 0.13 | -0.22 | 0.48 | |
| K | 0.20 | 0.06 | 0.34 | * |
| P | 0.32 | -1.02 | 1.66 | |
| Ph | -6.03 | -12.50 | 0.48 | |
| TKN | -0.01 | -0.02 | 0.00 | * |
| tmin.dry.mean | -0.70 | -3.27 | 1.89 | |
| pr.dry.mean | -0.03 | -0.08 | 0.03 | |
| tmax.dry.mean | 0.85 | -1.13 | 2.82 | |
| Ongokea gore | | | | |
| intercept | 0.01 | -0.84 | 0.87 | |
| hunting1 | 2.70 | -8.11 | 13.40 | |
| logging1 | 2.97 | -6.62 | 12.50 | |
| Dist | 0.29 | -0.14 | 0.72 | |
| K | 0.07 | -0.09 | 0.23 | |
| P | 0.37 | -1.25 | 2.00 | |

| | | | | |
|---|---|---|---|---|
| Ph | -1.02 | -8.08 | 5.96 | |
| TKN | -0.01 | -0.02 | 0.00 | |
| tmin.dry.mean | 0.74 | -2.31 | 3.76 | |
| pr.dry.mean | -0.07 | -0.14 | -0.01 | * |
| tmax.dry.mean | -0.44 | -2.68 | 1.83 | |
| Pancovia laurentii | | | | |
| intercept | 0.16 | -0.60 | 0.91 | |
| hunting1 | 9.20 | -0.41 | 19.20 | |
| logging1 | 2.26 | -6.91 | 11.30 | |
| Dist | 0.54 | 0.13 | 0.98 | * |
| K | -0.01 | -0.16 | 0.14 | |
| P | 0.66 | -0.80 | 2.14 | |
| Ph | -3.30 | -10.20 | 3.54 | |
| TKN | -0.02 | -0.04 | 0.00 | * |
| tmin.dry.mean | -0.01 | -2.70 | 2.69 | |
| pr.dry.mean | 0.04 | -0.02 | 0.10 | |
| tmax.dry.mean | -0.16 | -2.24 | 1.90 | |
| Pancovia pedicellaris | | | | |
| intercept | -0.15 | -1.04 | 0.70 | |
| hunting1 | 0.68 | -10.70 | 11.90 | |
| logging1 | -1.02 | -10.20 | 7.97 | |
| Dist | 0.36 | -0.05 | 0.78 | |
| K | -0.07 | -0.22 | 0.09 | |
| P | 1.42 | -0.20 | 3.12 | |
| Ph | -2.91 | -10.30 | 4.26 | |
| TKN | 0.00 | -0.02 | 0.02 | |
| tmin.dry.mean | -0.97 | -3.78 | 1.83 | |
| pr.dry.mean | 0.00 | -0.06 | 0.06 | |
| tmax.dry.mean | 0.74 | -1.36 | 2.89 | |
| Panda oleosa | | | | |
| intercept | 0.33 | -0.39 | 1.03 | |
| hunting1 | 3.19 | -5.71 | 11.90 | |
| logging1 | -3.64 | -11.80 | 4.55 | |
| Dist | -0.11 | -0.46 | 0.24 | |
| K | -0.12 | -0.26 | 0.02 | |
| P | 0.95 | -0.38 | 2.30 | |
| Ph | 2.27 | -4.09 | 8.64 | |
| TKN | 0.00 | -0.01 | 0.01 | |
| tmin.dry.mean | 1.06 | -1.52 | 3.66 | |

| | | | | |
|---|---|---|---|---|
| pr.dry.mean | -0.01 | -0.06 | 0.04 | |
| tmax.dry.mean | -0.40 | -2.36 | 1.56 | |
| Pausinystalia macroceras | | | | |
| intercept | 0.67 | -0.08 | 1.42 | |
| hunting1 | 5.20 | -3.89 | 14.40 | |
| logging1 | -1.59 | -9.93 | 6.70 | |
| Dist | 0.29 | -0.08 | 0.66 | |
| K | -0.07 | -0.22 | 0.07 | |
| P | -0.03 | -1.40 | 1.35 | |
| Ph | 5.00 | -1.85 | 11.90 | |
| TKN | -0.01 | -0.03 | 0.00 | * |
| tmin.dry.mean | -1.32 | -3.89 | 1.25 | |
| pr.dry.mean | 0.03 | -0.02 | 0.08 | |
| tmax.dry.mean | 0.51 | -1.48 | 2.51 | |
| Petersianthus macrocarpus | | | | |
| intercept | -0.08 | -0.78 | 0.63 | |
| hunting1 | 9.89 | 1.13 | 18.80 | * |
| logging1 | 1.83 | -6.59 | 10.20 | |
| Dist | 0.24 | -0.10 | 0.59 | |
| K | 0.14 | 0.00 | 0.28 | |
| P | 1.06 | -0.26 | 2.40 | |
| Ph | -6.30 | -12.60 | 0.08 | |
| TKN | -0.02 | -0.03 | -0.01 | * |
| tmin.dry.mean | 1.65 | -0.89 | 4.19 | |
| pr.dry.mean | -0.03 | -0.08 | 0.02 | |
| tmax.dry.mean | -0.70 | -2.65 | 1.24 | |
| Phyllocosmus africanus | | | | |
| intercept | 0.22 | -0.59 | 1.03 | |
| hunting1 | -1.32 | -11.60 | 8.49 | |
| logging1 | -2.42 | -11.20 | 6.31 | |
| Dist | -0.17 | -0.59 | 0.24 | |
| K | -0.04 | -0.21 | 0.12 | |
| P | -0.64 | -2.16 | 0.85 | |
| Ph | 2.54 | -4.52 | 9.63 | |
| TKN | 0.00 | -0.01 | 0.01 | |
| tmin.dry.mean | -2.45 | -5.71 | 0.75 | |
| pr.dry.mean | 0.01 | -0.05 | 0.07 | |
| tmax.dry.mean | 1.85 | -0.54 | 4.30 | |
| Picralima nitida | | | | |

| | | | | |
|---|---|---|---|---|
| intercept | -0.06 | -0.90 | 0.77 | |
| hunting1 | -4.31 | -15.80 | 6.60 | |
| logging1 | -1.13 | -10.30 | 7.98 | |
| Dist | -0.31 | -0.75 | 0.13 | |
| K | -0.09 | -0.26 | 0.07 | |
| P | 0.11 | -1.47 | 1.72 | |
| Ph | 1.63 | -5.54 | 8.76 | |
| TKN | 0.00 | -0.01 | 0.01 | |
| tmin.dry.mean | 0.34 | -3.14 | 3.80 | |
| pr.dry.mean | 0.00 | -0.07 | 0.06 | |
| tmax.dry.mean | 0.28 | -2.21 | 2.83 | |
| Pteleopsis hylodendron | | | | |
| intercept | -0.24 | -1.10 | 0.60 | |
| hunting1 | 0.33 | -9.79 | 10.40 | |
| logging1 | -5.10 | -14.20 | 3.89 | |
| Dist | -0.13 | -0.53 | 0.26 | |
| K | -0.02 | -0.19 | 0.15 | |
| P | 0.47 | -1.11 | 2.12 | |
| Ph | -3.68 | -11.70 | 3.88 | |
| TKN | 0.00 | -0.01 | 0.01 | |
| tmin.dry.mean | 0.36 | -2.39 | 3.18 | |
| pr.dry.mean | 0.02 | -0.04 | 0.07 | |
| tmax.dry.mean | 0.40 | -1.82 | 2.62 | |
| Pterocarpus soyauxii | | | | |
| intercept | -0.24 | -0.95 | 0.47 | |
| hunting1 | -0.01 | -8.96 | 8.82 | |
| logging1 | 0.96 | -7.18 | 9.09 | |
| Dist | 0.00 | -0.36 | 0.35 | |
| K | 0.03 | -0.12 | 0.17 | |
| P | 0.01 | -1.33 | 1.34 | |
| Ph | -3.11 | -9.47 | 3.25 | |
| TKN | 0.00 | -0.01 | 0.01 | |
| tmin.dry.mean | -0.03 | -2.60 | 2.57 | |
| pr.dry.mean | 0.00 | -0.05 | 0.05 | |
| tmax.dry.mean | 0.48 | -1.47 | 2.46 | |
| Radlkofera calodendron | | | | |
| intercept | -0.19 | -0.99 | 0.60 | |
| hunting1 | 1.85 | -7.91 | 11.80 | |
| logging1 | -9.16 | -18.20 | -0.30 | * |

| | | | | |
|---|---|---|---|---|
| Dist | -0.12 | -0.50 | 0.26 | |
| K | -0.11 | -0.29 | 0.05 | |
| P | 0.77 | -0.77 | 2.36 | |
| Ph | -4.57 | -12.10 | 2.70 | |
| TKN | -0.01 | -0.02 | 0.01 | |
| tmin.dry.mean | 0.22 | -2.46 | 2.90 | |
| pr.dry.mean | 0.00 | -0.05 | 0.06 | |
| tmax.dry.mean | 1.27 | -0.92 | 3.48 | |
| Rinorea oblongifolia | | | | |
| intercept | -0.91 | -1.91 | 0.16 | |
| hunting1 | -27.60 | -36.40 | -15.40 | * |
| logging1 | -14.30 | -28.20 | -2.40 | * |
| Dist | -1.03 | -1.57 | -0.49 | * |
| K | -0.13 | -0.43 | 0.14 | |
| P | -2.09 | -4.22 | 0.10 | |
| Ph | 3.47 | -7.35 | 14.50 | |
| TKN | 0.01 | -0.01 | 0.03 | |
| tmin.dry.mean | 0.32 | -4.11 | 4.75 | |
| pr.dry.mean | 0.07 | -0.02 | 0.16 | |
| tmax.dry.mean | 1.27 | -1.91 | 4.59 | |
| Santiria trimera | | | | |
| intercept | 0.12 | -0.63 | 0.87 | |
| hunting1 | 5.58 | -3.51 | 14.80 | |
| logging1 | 1.49 | -7.36 | 10.30 | |
| Dist | 0.32 | -0.05 | 0.70 | |
| K | 0.08 | -0.07 | 0.23 | |
| P | -0.09 | -1.58 | 1.40 | |
| Ph | -1.36 | -8.12 | 5.41 | |
| TKN | 0.00 | -0.01 | 0.01 | |
| tmin.dry.mean | 0.48 | -2.22 | 3.20 | |
| pr.dry.mean | 0.01 | -0.05 | 0.06 | |
| tmax.dry.mean | -0.83 | -2.97 | 1.26 | |
| Scottellia klaineana | | | | |
| intercept | 0.28 | -0.50 | 1.06 | |
| hunting1 | 3.24 | -5.94 | 12.50 | |
| logging1 | -1.70 | -10.60 | 7.19 | |
| Dist | 0.31 | -0.07 | 0.71 | |
| K | 0.00 | -0.16 | 0.16 | |
| P | -0.52 | -1.98 | 0.96 | |

| | | | | |
|---|---|---|---|---|
| Ph | 1.09 | -6.40 | 8.49 | |
| TKN | -0.01 | -0.02 | 0.01 | |
| tmin.dry.mean | -1.64 | -4.30 | 1.03 | |
| pr.dry.mean | 0.02 | -0.04 | 0.08 | |
| tmax.dry.mean | 0.81 | -1.25 | 2.91 | |
| Sterculia oblonga | | | | |
| intercept | 0.46 | -0.27 | 1.20 | |
| hunting1 | 2.85 | -6.18 | 11.90 | |
| logging1 | -6.87 | -15.80 | 1.96 | |
| Dist | -0.19 | -0.56 | 0.17 | |
| K | -0.08 | -0.22 | 0.06 | |
| P | 1.34 | -0.04 | 2.72 | |
| Ph | 3.57 | -2.97 | 10.10 | |
| TKN | -0.01 | -0.02 | 0.01 | |
| tmin.dry.mean | -0.18 | -2.99 | 2.60 | |
| pr.dry.mean | -0.02 | -0.08 | 0.04 | |
| tmax.dry.mean | 0.32 | -1.81 | 2.45 | |
| Strombosia grandifolia | | | | |
| intercept | -0.22 | -1.06 | 0.61 | |
| hunting1 | -0.44 | -10.50 | 9.37 | |
| logging1 | 1.40 | -7.17 | 10.10 | |
| Dist | 0.13 | -0.26 | 0.51 | |
| K | 0.04 | -0.11 | 0.18 | |
| P | 0.43 | -0.92 | 1.79 | |
| Ph | -2.74 | -10.20 | 4.63 | |
| TKN | 0.01 | 0.00 | 0.02 | |
| tmin.dry.mean | -0.76 | -3.49 | 1.97 | |
| pr.dry.mean | -0.03 | -0.08 | 0.03 | |
| tmax.dry.mean | 0.55 | -1.50 | 2.62 | |
| Strombosia nigropunctata | | | | |
| intercept | 1.77 | 0.81 | 2.77 | * |
| hunting1 | 13.00 | 1.89 | 24.70 | * |
| logging1 | -5.00 | -15.00 | 5.49 | |
| Dist | 0.53 | 0.09 | 1.01 | * |
| K | -0.22 | -0.43 | 0.00 | * |
| P | 2.18 | 0.20 | 4.24 | * |
| Ph | 14.40 | 4.80 | 24.20 | * |
| TKN | -0.01 | -0.03 | 0.01 | |
| tmin.dry.mean | -1.69 | -5.62 | 2.26 | |

| | | | |
|---|---|---|---|
| pr.dry.mean | 0.01 | -0.07 | 0.09 |
| tmax.dry.mean | -0.41 | -3.43 | 2.60 |
| Strombosia pustulata | | | |
| intercept | 0.44 | -0.28 | 1.16 |
| hunting1 | 3.74 | -5.09 | 12.50 |
| logging1 | 0.07 | -8.14 | 8.25 |
| Dist | 0.17 | -0.19 | 0.52 |
| K | -0.09 | -0.22 | 0.05 |
| P | 0.30 | -1.04 | 1.64 |
| Ph | 2.95 | -3.57 | 9.46 |
| TKN | 0.00 | -0.01 | 0.01 |
| tmin.dry.mean | -1.25 | -3.85 | 1.30 |
| pr.dry.mean | 0.01 | -0.04 | 0.06 |
| tmax.dry.mean | 0.70 | -1.28 | 2.67 |
| Strombosiopsis tetrandra | | | |
| intercept | 0.55 | -0.16 | 1.27 |
| hunting1 | 4.30 | -4.56 | 13.20 |
| logging1 | 0.36 | -7.84 | 8.53 |
| Dist | 0.19 | -0.16 | 0.55 |
| K | -0.10 | -0.24 | 0.05 |
| P | 0.99 | -0.35 | 2.34 |
| Ph | 4.93 | -1.50 | 11.40 |
| TKN | 0.00 | -0.01 | 0.01 |
| tmin.dry.mean | 0.73 | -1.82 | 3.28 |
| pr.dry.mean | -0.01 | -0.06 | 0.04 |
| tmax.dry.mean | -1.01 | -2.96 | 0.92 |
| Synsepalum longecuneatum | | | |
| intercept | 0.52 | -0.29 | 1.36 |
| hunting1 | 4.88 | -4.90 | 14.70 |
| logging1 | -2.49 | -11.70 | 6.42 |
| Dist | 0.22 | -0.17 | 0.61 |
| K | -0.10 | -0.26 | 0.06 |
| P | 1.34 | -0.15 | 2.87 |
| Ph | 3.80 | -3.37 | 11.30 |
| TKN | 0.00 | -0.02 | 0.01 |
| tmin.dry.mean | 0.16 | -2.44 | 2.79 |
| pr.dry.mean | 0.01 | -0.05 | 0.06 |
| tmax.dry.mean | -0.77 | -2.79 | 1.25 |
| Tabernaemontana penduliflora | | | |

| | | | | |
|---|---|---|---|---|
| intercept | -0.58 | -1.48 | 0.27 | |
| hunting1 | -3.33 | -13.50 | 6.63 | |
| logging1 | 2.16 | -7.42 | 12.00 | |
| Dist | -0.03 | -0.43 | 0.37 | |
| K | -0.02 | -0.20 | 0.14 | |
| P | -0.34 | -1.87 | 1.14 | |
| Ph | -5.79 | -13.70 | 1.76 | |
| TKN | -0.01 | -0.02 | 0.01 | |
| tmin.dry.mean | 0.04 | -2.92 | 3.02 | |
| pr.dry.mean | -0.03 | -0.09 | 0.03 | |
| tmax.dry.mean | 1.34 | -0.92 | 3.62 | |
| Terminalia superba | | | | |
| intercept | 0.13 | -0.61 | 0.89 | |
| hunting1 | -3.34 | -12.30 | 5.64 | |
| logging1 | -5.56 | -13.80 | 2.56 | |
| Dist | -0.25 | -0.60 | 0.11 | |
| K | 0.01 | -0.13 | 0.14 | |
| P | -0.35 | -1.67 | 0.96 | |
| Ph | 2.53 | -4.06 | 9.22 | |
| TKN | -0.01 | -0.02 | 0.00 | |
| tmin.dry.mean | -1.90 | -4.63 | 0.77 | |
| pr.dry.mean | 0.00 | -0.05 | 0.06 | |
| tmax.dry.mean | 1.67 | -0.42 | 3.79 | |
| Thomandersia hensii | | | | |
| intercept | 0.85 | 0.02 | 1.72 | * |
| hunting1 | 3.58 | -6.21 | 13.40 | |
| logging1 | -4.77 | -13.30 | 3.65 | |
| Dist | 0.18 | -0.21 | 0.57 | |
| K | -0.12 | -0.28 | 0.04 | |
| P | 0.59 | -0.81 | 2.00 | |
| Ph | 8.40 | 0.82 | 16.30 | * |
| TKN | -0.01 | -0.02 | 0.01 | |
| tmin.dry.mean | -0.79 | -3.50 | 1.91 | |
| pr.dry.mean | 0.00 | -0.06 | 0.06 | |
| tmax.dry.mean | -0.21 | -2.33 | 1.90 | |
| Trichilia prieuriana | | | | |
| intercept | 0.27 | -0.43 | 0.98 | |
| hunting1 | -0.31 | -8.93 | 8.45 | |
| logging1 | 0.15 | -7.99 | 8.17 | |

| | | | | |
|---|---|---|---|---|
| Dist | -0.08 | -0.42 | 0.27 | |
| K | -0.12 | -0.26 | 0.01 | |
| P | 0.78 | -0.54 | 2.10 | |
| Ph | 3.11 | -3.24 | 9.47 | |
| TKN | 0.01 | 0.00 | 0.02 | |
| tmin.dry.mean | -1.24 | -3.74 | 1.28 | |
| pr.dry.mean | 0.02 | -0.03 | 0.07 | |
| tmax.dry.mean | 0.75 | -1.18 | 2.69 | |
| Trichilia rubescens | | | | |
| intercept | 0.25 | -0.47 | 0.97 | |
| hunting1 | -4.01 | -12.90 | 4.91 | |
| logging1 | -3.45 | -11.80 | 4.89 | |
| Dist | -0.32 | -0.67 | 0.03 | |
| K | 0.09 | -0.05 | 0.23 | |
| P | -0.11 | -1.49 | 1.31 | |
| Ph | 6.00 | -0.39 | 12.40 | |
| TKN | 0.02 | 0.01 | 0.03 | * |
| tmin.dry.mean | 1.00 | -1.53 | 3.54 | |
| pr.dry.mean | -0.02 | -0.08 | 0.03 | |
| tmax.dry.mean | -1.51 | -3.49 | 0.44 | |
| Trichilia welwitschii | | | | |
| intercept | 0.36 | -0.42 | 1.17 | |
| hunting1 | 0.48 | -8.76 | 9.77 | |
| logging1 | -2.38 | -10.70 | 5.75 | |
| Dist | -0.21 | -0.57 | 0.14 | |
| K | 0.07 | -0.07 | 0.20 | |
| P | 0.28 | -1.06 | 1.61 | |
| Ph | 4.93 | -1.98 | 12.00 | |
| TKN | 0.00 | -0.01 | 0.01 | |
| tmin.dry.mean | 1.86 | -0.85 | 4.52 | |
| pr.dry.mean | -0.08 | -0.13 | -0.02 | * |
| tmax.dry.mean | -1.41 | -3.51 | 0.70 | |
| Vitex welwitschii | | | | |
| intercept | 0.25 | -0.55 | 1.04 | |
| hunting1 | 5.25 | -5.69 | 16.00 | |
| logging1 | -2.97 | -13.60 | 7.90 | |
| Dist | -0.03 | -0.46 | 0.39 | |
| K | 0.01 | -0.15 | 0.16 | |
| P | 0.63 | -1.05 | 2.32 | |

| | | | | |
|---|---|---|---|---|
| Ph | -0.12 | -7.09 | 6.93 | |
| TKN | -0.01 | -0.02 | 0.01 | |
| tmin.dry.mean | -0.12 | -3.13 | 2.89 | |
| pr.dry.mean | 0.03 | -0.04 | 0.09 | |
| tmax.dry.mean | -0.12 | -2.33 | 2.09 | |
| Xylopia chrysophylla | | | | |
| intercept | 0.11 | -0.70 | 0.91 | |
| hunting1 | 6.39 | -4.30 | 17.10 | |
| logging1 | 4.61 | -5.77 | 16.10 | |
| Dist | 0.27 | -0.17 | 0.74 | |
| K | 0.02 | -0.13 | 0.17 | |
| P | -0.65 | -2.38 | 1.05 | |
| Ph | -2.33 | -9.90 | 4.92 | |
| TKN | 0.00 | -0.02 | 0.01 | |
| tmin.dry.mean | -0.86 | -3.82 | 2.01 | |
| pr.dry.mean | 0.04 | -0.02 | 0.10 | |
| tmax.dry.mean | 0.38 | -1.76 | 2.57 | |
| Xylopia phloiodora | | | | |
| intercept | 0.52 | -0.26 | 1.33 | |
| hunting1 | 5.23 | -4.91 | 15.60 | |
| logging1 | 0.19 | -8.33 | 8.89 | |
| Dist | 0.27 | -0.13 | 0.68 | |
| K | -0.09 | -0.25 | 0.06 | |
| P | -0.53 | -2.08 | 1.00 | |
| Ph | 3.40 | -3.68 | 10.60 | |
| TKN | -0.01 | -0.02 | 0.01 | |
| tmin.dry.mean | -0.49 | -3.29 | 2.32 | |
| pr.dry.mean | 0.01 | -0.05 | 0.07 | |
| tmax.dry.mean | 0.18 | -2.01 | 2.38 | |
| Zanthoxylum gilletii | | | | |
| intercept | 0.00 | -0.90 | 0.91 | |
| hunting1 | -11.50 | -23.30 | 0.37 | |
| logging1 | -11.60 | -21.80 | -1.70 | * |
| Dist | -0.75 | -1.25 | -0.24 | * |
| K | -0.09 | -0.27 | 0.08 | |
| P | -0.14 | -1.76 | 1.47 | |
| Ph | 6.45 | -1.05 | 14.00 | |
| TKN | -0.01 | -0.03 | 0.00 | * |
| tmin.dry.mean | 1.67 | -2.24 | 5.62 | |

| | | | |
|---|---|---|---|
| pr.dry.mean | 0.00 | -0.08 | 0.07 |
| tmax.dry.mean | -0.15 | -2.92 | 2.66 |

**A3: Table of community sensitivity to environmental covariates. Sensitivity is dimensionless and on the scale of the predictor.**

Community Sensitivities to Predictors

| Predictor | Estimate | 2.5% | 97.5% |
|---|---|---|---|
| Dry Season Precip. | 0.916 | 0.756 | 1.09 |
| Soil pH | 1.02 | 0.847 | 1.2 |
| Soil K | 1.08 | 0.908 | 1.27 |
| Soil P | 1.11 | 0.922 | 1.31 |
| Soil TKN | 1.15 | 0.94 | 1.38 |
| Dry Season Max. Temp. | 1.28 | 1.06 | 1.51 |
| Dry Season Min. Temp. | 1.48 | 1.21 | 1.78 |
| Logging | 1.99 | 1.65 | 2.35 |
| Hunting | 2.22 | 1.85 | 2.61 |
| Distance to Village | 3.29 | 2.76 | 3.85 |

### A4: Trait-specific posterior parameter estimates for the effect of environmental covariates on community weighted trait values.

| Posterior Parameter Estimates for Traits | | | | |
|---|---|---|---|---|
| | **Estimate** | **2.50%** | **97.50%** | **0 ∉ 95% CI** |
| **Leaf Toughness** | | | | |
| Intercept | 0.0147 | 0.00267 | 0.0269 | * |
| Hunting | 0.133 | -0.0169 | 0.283 | |
| Logging | 0.00459 | -0.136 | 0.143 | |
| Dist. to Village | 0.00147 | -0.00448 | 0.00739 | |
| Soil K | -0.000807 | -0.00314 | 0.00151 | |
| Soil P | 0.0176 | -0.00492 | 0.0404 | |
| Soil Ph | 0.0957 | -0.0133 | 0.206 | |
| Soil TKN | 1.33E-05 | -0.000162 | 0.000188 | |
| Dry Min. Temp. | -0.000618 | -0.0437 | 0.0425 | |
| Dry Precip. | 0.000119 | -0.00077 | 0.001 | |
| Dry Max. Temp. | 0.0366 | 0.00348 | 0.0697 | * |
| **Sapwood Density** | | | | |
| Intercept | 0.00661 | 0.00386 | 0.00933 | * |
| Hunting | 0.0326 | -0.00125 | 0.0664 | |
| Logging | -0.000544 | -0.032 | 0.0312 | |
| Dist. to Village | 0.00141 | 7.34E-05 | 0.00276 | * |
| Soil K | -0.000454 | -0.00098 | 7.07E-05 | |
| Soil P | 0.00169 | -0.00338 | 0.00679 | |
| Soil Ph | 0.0575 | 0.0326 | 0.0825 | * |
| Soil TKN | 6.19E-06 | -3.33E-05 | 4.59E-05 | |
| Dry Min. Temp. | -0.00866 | -0.0185 | 0.00117 | |
| Dry Precip. | 0.000138 | -6.22E-05 | 0.000341 | |
| Dry Max. Temp. | 0.0176 | 0.0101 | 0.0251 | * |
| **Leaf N** | | | | |
| Intercept | 0.000194 | 0.00012 | 0.000268 | * |
| Hunting | 0.00117 | 0.000259 | 0.00208 | * |
| Logging | 9.16E-05 | -0.000762 | 0.000939 | |
| Dist. to Village | 7.25E-06 | -2.91E-05 | 4.35E-05 | |
| Soil K | -1.26E-05 | -2.67E-05 | 1.61E-06 | |
| Soil P | 6.18E-05 | -7.62E-05 | 0.000199 | |
| Soil Ph | 0.00149 | 0.000827 | 0.00217 | * |
| Soil TKN | 1.35E-06 | 2.70E-07 | 2.41E-06 | * |
| Dry Min. Temp. | -0.00032 | -0.000584 | -5.24E-05 | * |
| Dry Precip. | 3.82E-06 | -1.67E-06 | 9.31E-06 | |
| Dry Max. Temp. | 0.000698 | 0.000493 | 0.000899 | * |
| **Leaf C:N ratio** | | | | |
| Intercept | 0.22 | 0.125 | 0.315 | * |
| Hunting | 0.869 | -0.309 | 2.05 | |
| Logging | -0.335 | -1.44 | 0.771 | |
| Dist. to Village | 0.0343 | -0.0126 | 0.0814 | |
| Soil K | -0.0169 | -0.0353 | 0.00149 | |
| Soil P | 0.202 | 0.0238 | 0.381 | * |
| Soil Ph | 2.06 | 1.19 | 2.92 | * |
| Soil TKN | 0.000243 | -0.00114 | 0.00164 | |
| Dry Min. Temp. | -0.102 | -0.445 | 0.241 | |
| Dry Precip. | 0.000929 | -0.00618 | 0.008 | |
| Dry Max. Temp. | 0.516 | 0.256 | 0.779 | * |
| **Leaf N15** | | | | |
| Intercept | 0.0239 | -0.0113 | 0.0593 | |
| Hunting | -0.0757 | -0.514 | 0.362 | |
| Logging | -0.00434 | -0.414 | 0.405 | |
| Dist. to Village | -0.0101 | -0.0276 | 0.00728 | |
| Soil K | -0.00185 | -0.00869 | 0.00498 | |
| Soil P | -0.00147 | -0.0676 | 0.0646 | |
| Soil Ph | 0.321 | 0.000991 | 0.64 | * |
| Soil TKN | 0.00061 | 9.62E-05 | 0.00112 | * |
| Dry Min. Temp. | -0.0398 | -0.168 | 0.0879 | |
| Dry Precip. | 0.000148 | -0.00245 | 0.00275 | |
| Dry Max. Temp. | 0.0644 | -0.0329 | 0.162 | |

| Posterior Parameter Estimates for Traits | | | | |
|---|---|---|---|---|
| | **Estimate** | **2.50%** | **97.50%** | **0 ∉ 95% CI** |
| **Leaf C13** | | | | |
| Intercept | -0.29 | -0.323 | -0.258 | * |
| Hunting | -1.42 | -1.82 | -1.02 | * |
| Logging | 0.0921 | -0.285 | 0.469 | |
| Dist. to Village | -0.0325 | -0.0485 | -0.0166 | * |
| Soil K | 0.0191 | 0.0129 | 0.0254 | * |
| Soil P | -0.181 | -0.242 | -0.12 | * |
| Soil Ph | -2.51 | -2.8 | -2.21 | * |
| Soil TKN | -0.00127 | -0.00174 | -0.000804 | * |
| Dry Min. Temp. | 0.289 | 0.172 | 0.406 | * |
| Dry Precip. | -0.00325 | -0.00565 | -0.000855 | * |
| Dry Max. Temp. | -0.839 | -0.929 | -0.75 | * |
| **Leaf Chlorophyll Concentration** | | | | |
| Intercept | 0.66 | 0.427 | 0.893 | * |
| Hunting | 3.75 | 0.848 | 6.65 | * |
| Logging | 0.653 | -2.05 | 3.36 | |
| Dist. to Village | 0.16 | 0.0456 | 0.275 | * |
| Soil K | -0.0439 | -0.0889 | 0.000823 | |
| Soil P | 0.352 | -0.0842 | 0.789 | |
| Soil Ph | 5.4 | 3.3 | 7.51 | * |
| Soil TKN | 0.00109 | -0.0023 | 0.00447 | |
| Dry Min. Temp. | -0.791 | -1.63 | 0.047 | |
| Dry Precip. | 0.0101 | -0.00711 | 0.0273 | |
| Dry Max. Temp. | 2 | 1.37 | 2.65 | * |
| **Leaf Surface Area** | | | | |
| Intercept | 0.0377 | 0.02 | 0.0557 | * |
| Hunting | 0.193 | -0.027 | 0.415 | |
| Logging | -0.0334 | -0.239 | 0.173 | |
| Dist. to Village | -0.00104 | -0.00979 | 0.00769 | |
| Soil K | -0.00257 | -0.006 | 0.000876 | |
| Soil P | 0.0346 | 0.00148 | 0.068 | * |
| Soil Ph | 0.318 | 0.157 | 0.482 | * |
| Soil TKN | 0.000164 | -9.54E-05 | 0.000422 | |
| Dry Min. Temp. | -0.029 | -0.0929 | 0.0352 | |
| Dry Precip. | 0.000106 | -0.00123 | 0.00142 | |
| Dry Max. Temp. | 0.112 | 0.0623 | 0.161 | * |
| **Leaf SLA** | | | | |
| Intercept | 0.0954 | 0.0236 | 0.167 | * |
| Hunting | 0.188 | -0.698 | 1.06 | |
| Logging | -0.154 | -0.988 | 0.669 | |
| Dist. to Village | 0.00507 | -0.0302 | 0.04 | |
| Soil K | -0.00855 | -0.0223 | 0.00519 | |
| Soil P | -0.0412 | -0.175 | 0.0916 | |
| Soil Ph | 0.893 | 0.252 | 1.54 | * |
| Soil TKN | 0.000354 | -0.000678 | 0.00139 | |
| Dry Min. Temp. | -0.269 | -0.525 | -0.0141 | * |
| Dry Precip. | 0.00348 | -0.00174 | 0.00876 | |
| Dry Max. Temp. | 0.479 | 0.283 | 0.676 | * |
| **Seed Mass** | | | | |
| Intercept | 0.0441 | 0.000899 | 0.0873 | * |
| Hunting | 0.176 | -0.355 | 0.709 | |
| Logging | 0.0879 | -0.415 | 0.59 | |
| Dist. to Village | 0.0116 | -0.0097 | 0.0328 | |
| Soil K | 5.19E-05 | -0.00835 | 0.00845 | |
| Soil P | 0.0276 | -0.0532 | 0.108 | |
| Soil Ph | 0.437 | 0.0458 | 0.83 | * |
| Soil TKN | 3.07E-05 | -0.000596 | 0.000661 | |
| Dry Min. Temp. | 0.00425 | -0.152 | 0.16 | |
| Dry Precip. | -0.00193 | -0.00514 | 0.00125 | |
| Dry Max. Temp. | 0.0695 | -0.0497 | 0.187 | |

**A5: Table of community weighted trait sensitivity to environmental covariates. Sensitivity is dimensionless and on the scale of the predictor.**

**Community Weighted Trait Sensitivities to Predictors**

| Predictor | Estimate | 2.50% | 97.50% |
|---|---|---|---|
| Dry Season Precipitation | 4720 | 198 | 13800 |
| Soil K | 6430 | 268 | 18400 |
| Dry Season Max. Temp | 6450 | 274 | 18900 |
| Soil TKN | 6650 | 289 | 18300 |
| Dry Season Min. Temp. | 8370 | 360 | 24300 |
| Soil pH | 8700 | 419 | 22100 |
| Soil P | 9920 | 499 | 25700 |
| Hunting | 10300 | 437 | 30200 |
| Logging | 10500 | 447 | 30200 |
| Distance from Village | 17900 | 793 | 48100 |

**A6: Trace of posterior chains for each environmental covariate shows convergence of parameter estimates.**

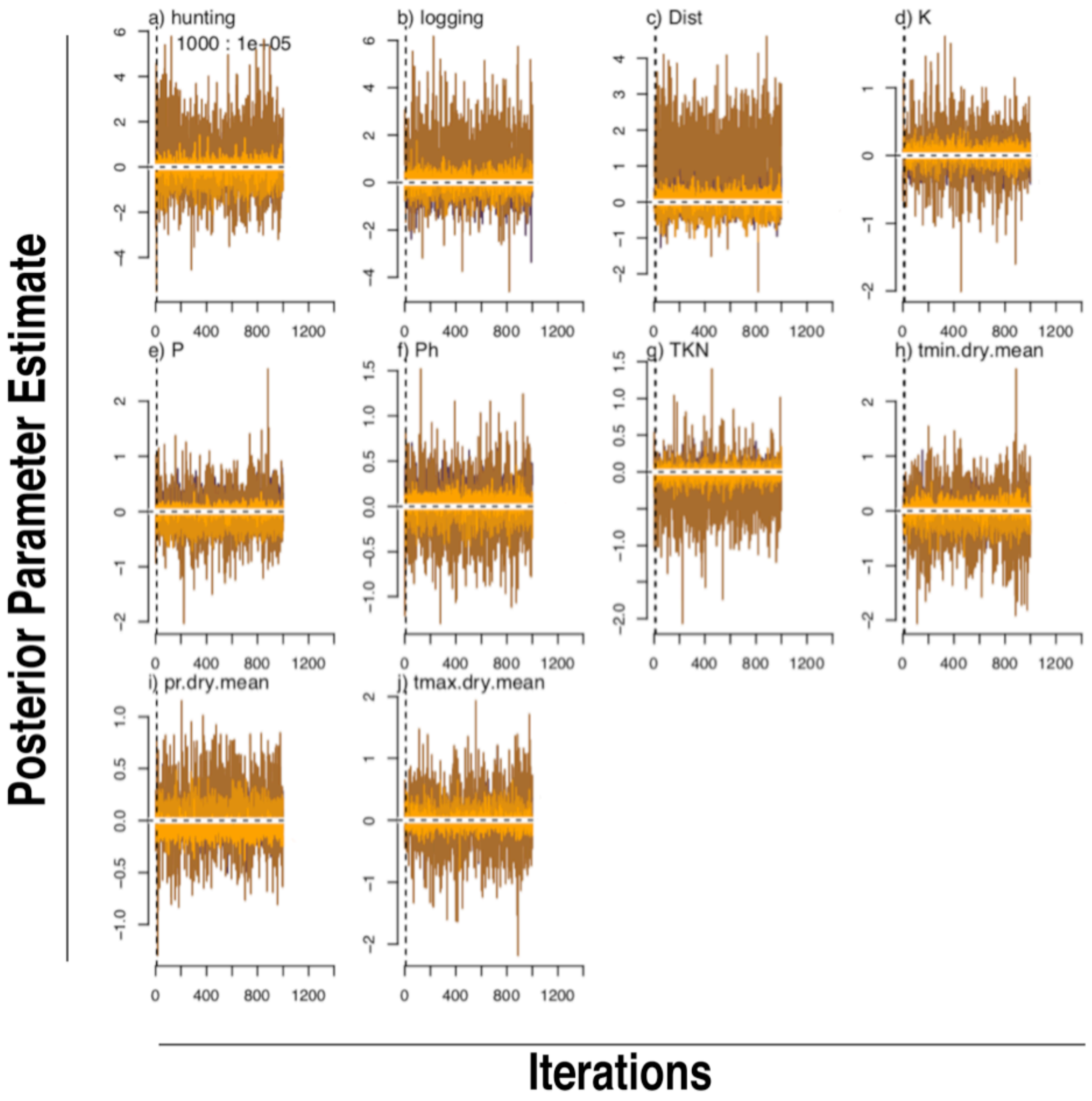

**A7: Community weighted trait ordination suggests that species are clustered into pioneer and secondary communities.**

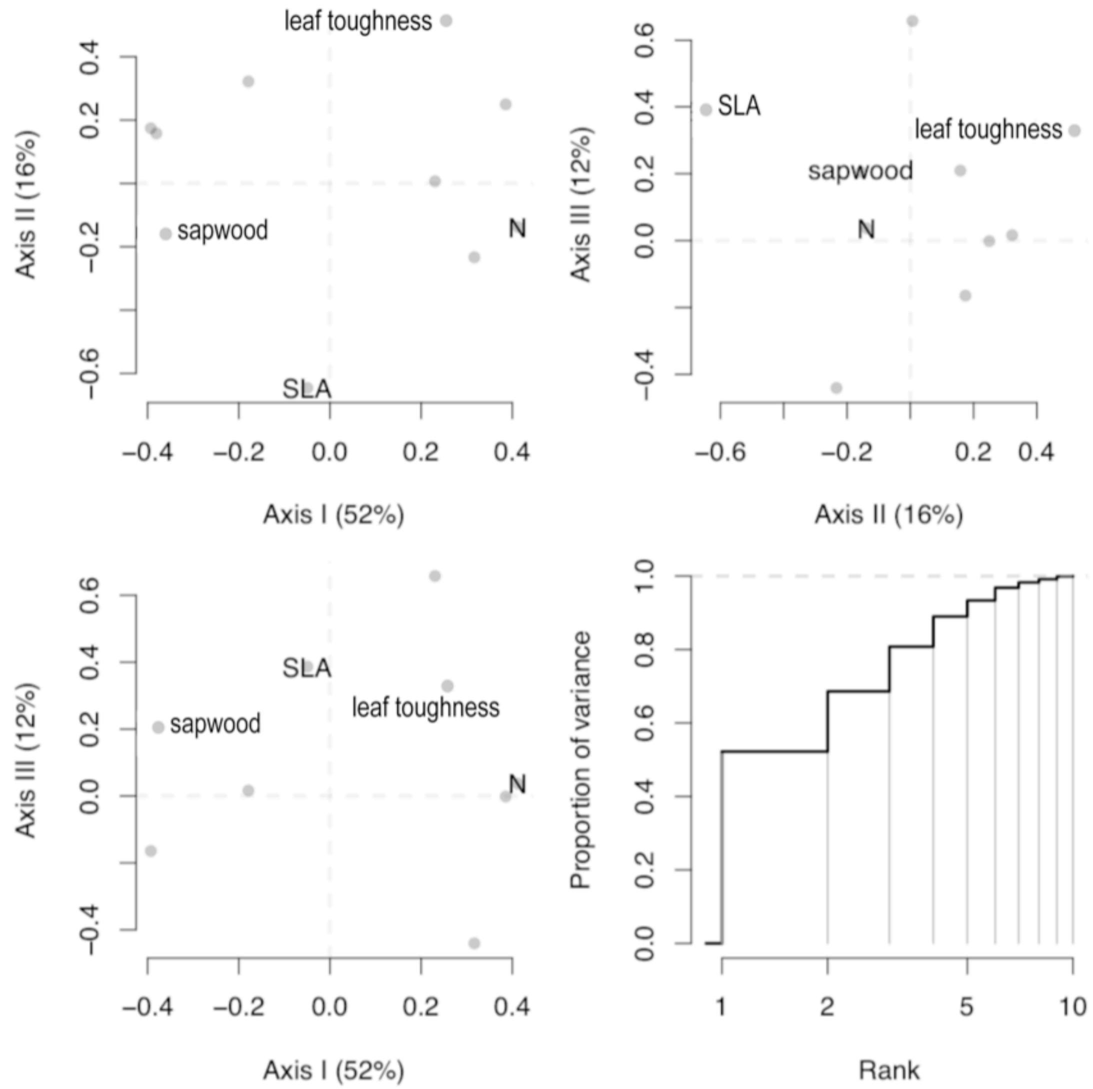

**A8: Dendrogram demonstrating clustering by correlation in trait data (A) and correlation by trait response to environment (B).**

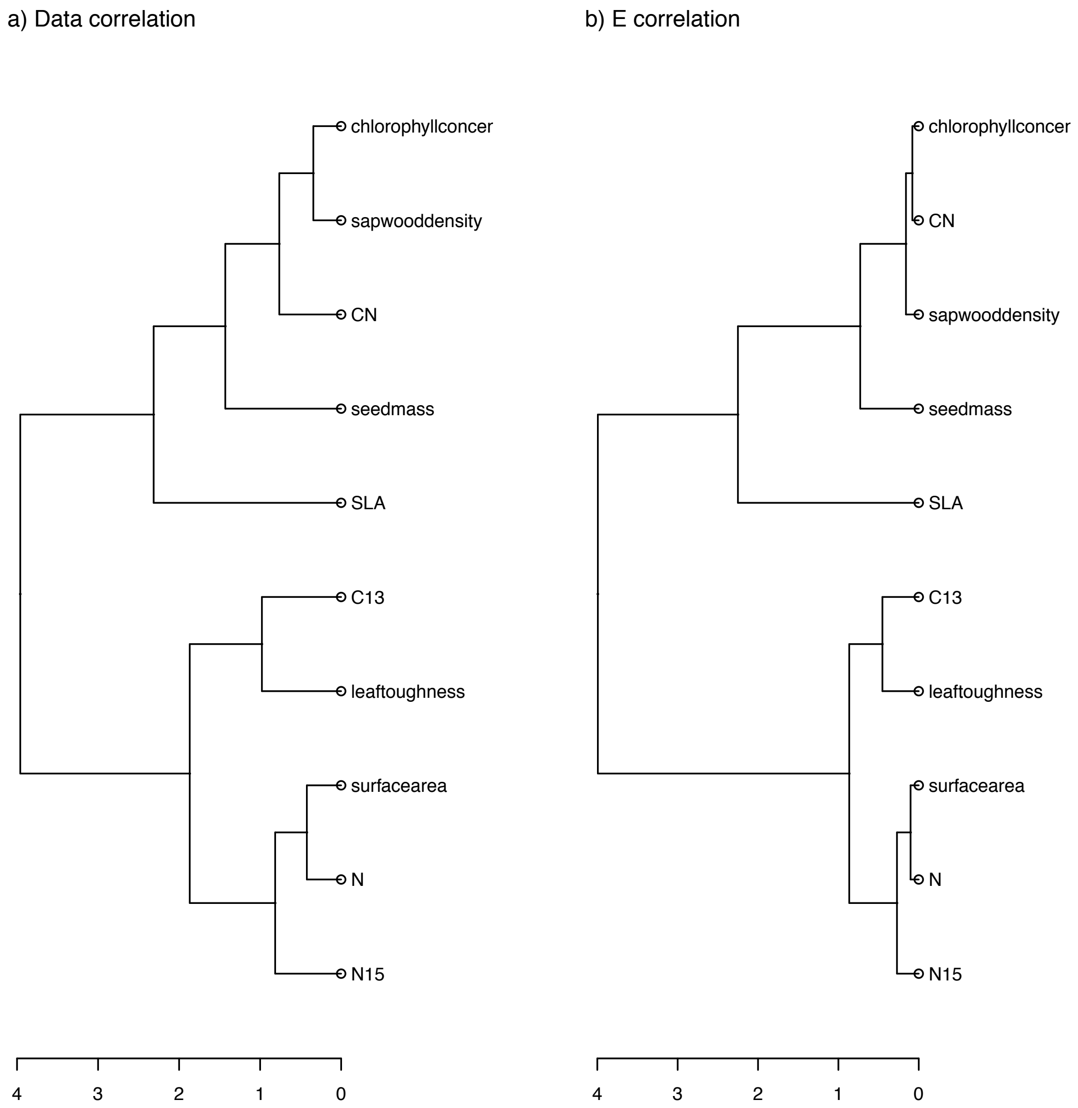

**A9: Ordered species names from figure 2 are clustered into a disturbance tolerant group (1), and a disturbance intolerant group (2).**

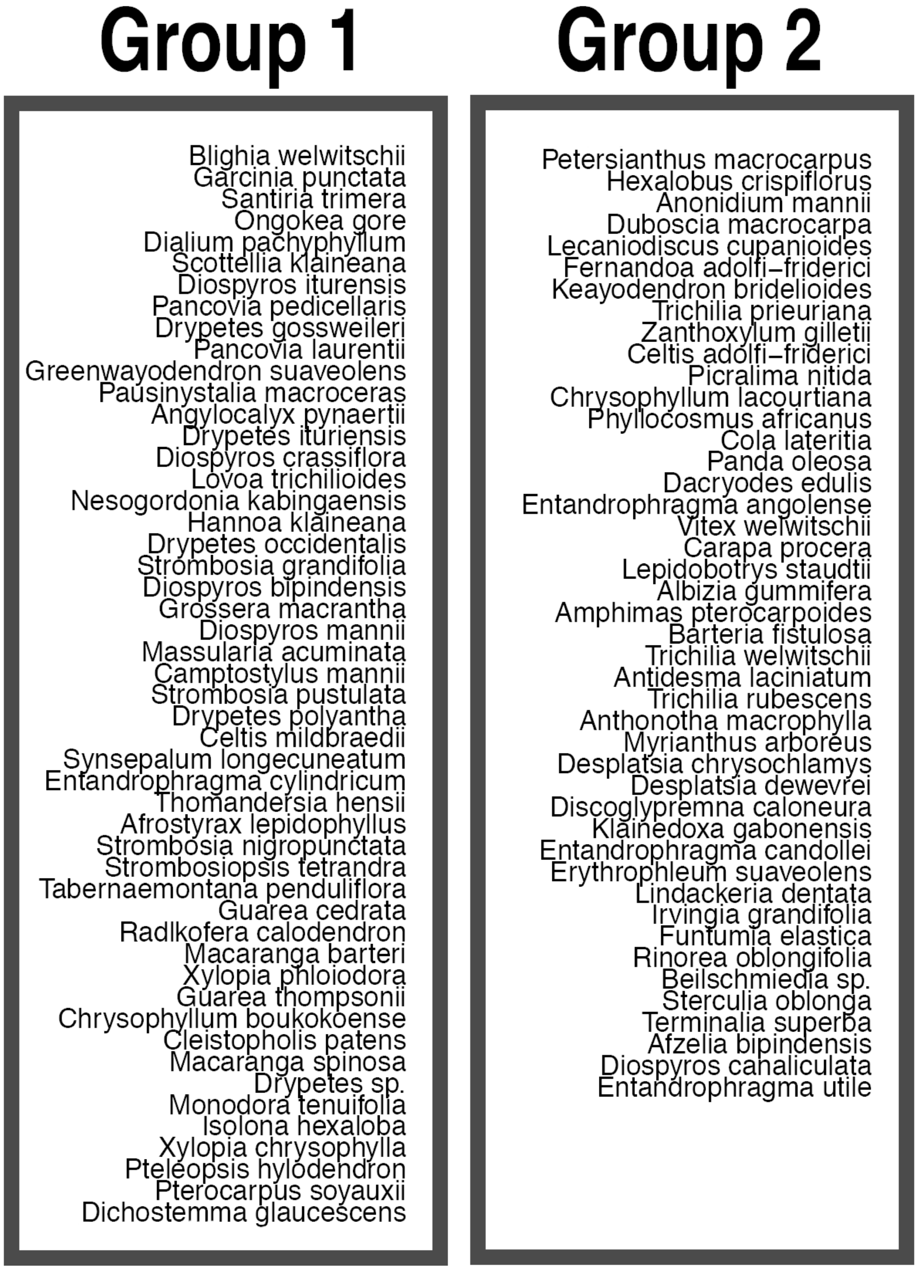
